# A mechanosensory feedback that uncouples external and self-generated sensory responses in the olfactory cortex

**DOI:** 10.1101/2023.04.04.535405

**Authors:** Alireza A. Dehaqani, Filippo Michelon, Paola Patella, Luigi Petrucco, Eugenio Piasini, Giuliano Iurilli

## Abstract

Sampling behaviors have sensory consequences that can hinder perceptual stability. In olfaction, sniffing affects early odor encoding, mimicking a sudden change in odor concentration. Here, we examined how the inhalation speed impacts the representation of odor concentration in the main olfactory cortex. Neurons combine the odor input with a global top-down signal preceding the sniff and a mechanosensory feedback generated by the air passage through the nose during an inhalation. Still, the population representation of concentration is remarkably sniff-invariant. This is because the mechanosensory and olfactory responses are uncorrelated within and across neurons. Thus, a faster odor inhalation and a concentration increase change the cortical activity pattern in distinct ways. This encoding strategy affords tolerance to potential concentration fluctuations caused by varying inhalation speeds. Since mechanosensory reafferences are widespread across sensory systems, the coding scheme described here may be a canonical strategy to mitigate the sensory ambiguities caused by movements.

## INTRODUCTION

A problem animals face as they explore the world is that their own actions generate sensory stimuli that can be indistinguishable from those caused by external events. A consequence of this ambiguity is that stimulus encoding becomes subject to the moment-by-moment variability of sampling movements^1^. In olfaction, this problem arises whenever an animal changes how fast it samples an odorant^2–5^. In terrestrial vertebrates, breathing through the nose provides the animal with repeated snapshots of the olfactory world^6^. Animals can compare these sequential samples to analyze the olfactory scene, detecting changes in odors and their concentration^7–9^. However, when an animal takes rapid breaths, the nasal airflow rate increases. As a result, the concentration profile of an odor inside the nasal cavity is thought to change, mimicking an increase in the environmental odor concentration^2–4^.

Odorants are detected by receptors expressed by the olfactory sensory neurons (OSNs). OSNs innervate the main olfactory bulbs (OBs), which transform olfactory information and broadcast their output to higher brain centers through projection neurons called mitral/tufted (MT) neurons. Consistent with the effect of the nasal airflow on the number of molecules that reach the olfactory epithelium, sniffing alters odor responses at the early stages of olfactory processing^10–12^. Remarkably, faster sniffs increase the magnitude and reduce the latency of the odor responses of the MT neurons as if the external concentration had suddenly increased^4,5^. Nevertheless, humans and rodents can quickly learn to distinguish the concentration of odors regardless of the inhalation speed^2,4,5^. How these physiological and behavioral results can be reconciled is a long-standing question.

It has been proposed that an animal can discriminate an increase in the environmental concentration from the effect of a faster inhalation because the olfactory system may encode how fast the animal is breathing^2–5^. This hypothesis is supported by the observation that nasal breathing entrains the activity of the olfactory system even in the absence of odors^11,13–19^; moreover, it has been demonstrated that the spiking activity of MT neurons encodes the duration of an inhalation^5^. Still, the origin and content of this information is unclear. Efferent copies of the inhalation command could predict the airflow kinematics in the nasal cavities; however, there is currently no evidence of motor inputs to the olfactory system. Instead, since many olfactory receptors respond to mechanical stimuli as well as to odorants^20,21^, the information about the inhalation speed could be supplied by the air passing through the nose during each inhalation. Notably, however, this latter possibility also highlights another potential problem arising with each inhalation. Because mechanosensory and olfactory inputs have the same source, the airflow signal could actually increase confusion and interfere with olfactory information, further hindering the representation of odor concentration. Thus, whether and how a non-olfactory signal could help distinguish the olfactory consequences of sniffing remains to be determined.

Here we set out to examine whether the speed of inhalation affects the representation of odor concentration in a higher-order area called piriform cortex (PCx). The PCx represents odors through a population code, in which the distributed activity of ensembles of neurons affords downstream brain areas systematic information about odor identity and concentration^22–28^. Recording the spiking activity of ensembles of neurons, we observed that odor responses are primarily sensitive to the mechanical stimulation of the olfactory epithelium by inhaled air and to a global top-down input that seems to precede sniffs. The mechanosensory response to a quick breath can increase or decrease relative to a regular inhalation, independent of how the neuron responds to a concentration increase. As a result, the mechanosensory response distinguishes the population representation of an odor during a sniff without interfering with the representation of the concentration of that odor, effectively ensuring a sniff-invariant concentration code.

## RESULTS

### Sniffing causes unstable odor responses in the PCx

To determine the effects of inhalation speed on odor responses in the PCx, we used Neuropixels 1.0 probes to record inhalation-by-inhalation responses to different odor concentrations in awake, head-fixed mice (Figure 1A-B). We monitored inhalations through a flow sensor in front of the mouse naris. The flow sensor measured the airflow rate at the entrance of the nasal cavity during each inhalation. Since the distribution of inhalation durations appeared bimodal, we used a semi-unsupervised algorithm that sorted inhalations into slow and fast depending on their airflow rate waveforms (Figure 1C and Figure S1A; see *Methods*). Fast inhalations had shorter duration and steeper changes in airflow rate than slow inhalations. We use “sniff” and “fast inhalation” interchangeably, regardless of whether the origin was autonomic or voluntary^29^. Our neural recordings sampled excitatory (semilunar and pyramidal) and inhibitory neurons from layers 2 and 3 of the anterior PCx. The early 400 ms after the onset of an odor presentation were dominated by quick sniffs (Figure S1B) and a typical transient response. As our goal was to investigate whether the representation of an environmental odor remained stable despite variable inhalation speed, we did not analyze the response during the early 400 ms of odor presentation, reducing other sources of variability. Similarly, we did not examine the odor responses during the first two trials of each odor-concentration stimulus to avoid variability due to the stimulus novelty.

**Figure 1.**
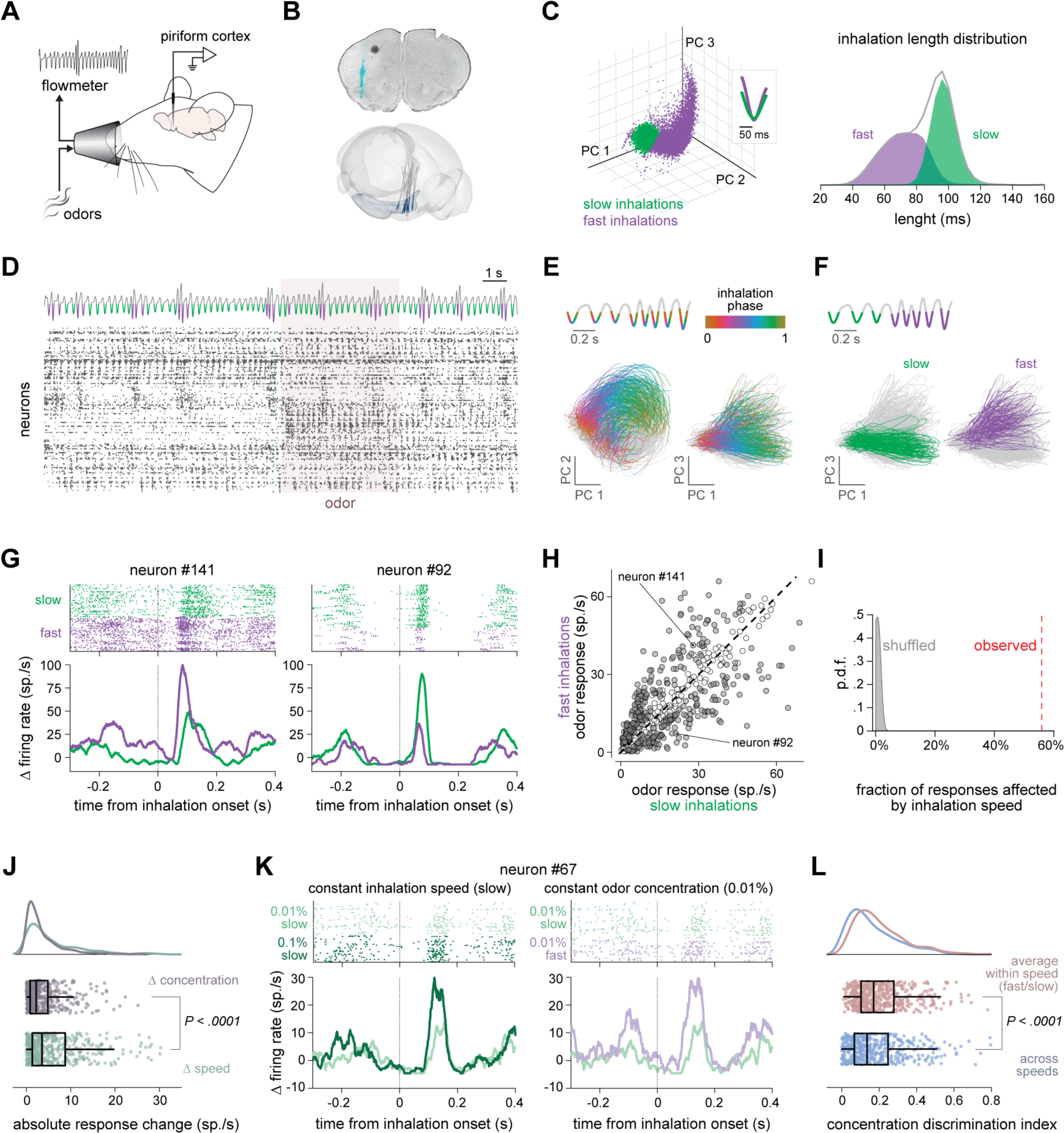
Sniffing causes unstable odor responses in the PCx. **(A)** Experimental setup for PCx recordings paired with odor delivering and inhalation monitoring. The speed of the airflow generated at the entrance of the naris by respiration was measured through a flowmeter. **(B)** Recording sites. Top, example of coronal section indicating the placement of a Neuropixels probe in the anterior PCx (cyan). Bottom, reconstructed locations of Neuropixels probes from all experiments (N = 22 mice). **(C)** Types of inhalations. Left, PCA plot of inhalation waveforms sorted via Gaussian-mixture model-hierarchical clustering; inset, average slow (green) and fast (purple) inhalation waveforms. Right, distribution of slow and fast inhalation lengths. Gray curve, length distribution before inhalation sorting. **(D)** Example of simultaneous PCx activity and respiratory airflow speed. External flow sensor signal (top) and spiking raster plot (bottom). Shaded area, odor period (5 seconds). Slow and fast inhalations are color-coded in green and purple, respectively. Neurons are sorted along the Y-axis of the raster plot based on their activity pattern to highlight the sequential activation of neurons with each inhalation. **(E)** PCA embedding of neural trajectories during individual respiratory cycles for an example mouse (time bin: 10 ms). The inhalation phase is color-coded according to the scheme in the inset plot on the top (inhalation onset: 0; offset: 1). Gray, exhalation. Inhalation phase decoding using the population activity is illustrated in Figure S1C. **(F)** Same neural trajectories as in (E), color-coded according to the inhalation speed. Green, slow inhalations; purple, fast inhalations. Inhalation speed decoding using the population activity is shown in Figure S1D. **(G)** Examples of odor responses during slow and fast inhalations. Raster plot (top) and PETH (bottom) for two neurons from the same mouse responding to the same odor and concentration. Responses to slow and fast inhalations are color-coded in purple and green, respectively. **(H)** Scatterplot of odor response amplitudes (average firing rate in 180 ms window and across slow or fast inhalations; five odors, 0.01 and 0.1% v./v. concentration; n = 687 odor-concentration responses) for all odor-responsive neurons (Benjamini-Hochberg adjusted *P*-value < 0.05, Wilcoxon signed rank test; n = 195 neurons, six mice). Gray dots, significant difference between the amplitudes of the responses during fast and slow inhalations of an odor presented at a given concentration; white dots, no statistical difference (*P*-value < 0.01, two-sided Wilcoxon rank-sum test). The odor responses of Figure 1G are indicated. **(I)** Fraction of odor concentration responses changing with inhalation speed (red vertical line) and null distribution obtained by shuffling 1000 times the inhalation speed label for each odor concentration response. **(J)** Absolute change in firing rate due to a 0.01-to-0.1% change in odor concentration and a slow-to-fast switch in inhalation of 0.01% odor (n = 442 neuron-odor pairs; only odor-responsive neurons; *P*-value < 0.0001, two-sided Wilcoxon rank-sum test). Top: probability density function of the two distributions. Bottom, boxplots indicate the median, 25th and 75th percentile (box edges), and 1.5 times the interquartile range (whiskers) of the two distributions. **(K)** Examples of responses from one neuron to different odor concentrations and inhalation speeds. Left, raster plot and PETHs of the responses to 0.01 % (light green) and 0.1 % (dark green) v./v. averaged across all inhalations. Right, raster plot and PETHs of the responses to slow (light green) and fast (light purple) inhalations of 0.01% v./v. odor. **(L)** CDI calculated using odor concentration responses within and across inhalation speeds (*P* - value < 0.0001, two-sided Wilcoxon rank-sum test). Within-inhalation-speed CDIs are averages of the only-slow and only-fast CDIs. Across-inhalation speed CDIs were calculated using all neuron responses regardless of the inhalation speed. Same neuron-odor pairs as in (J). Top, probability density function of the two distributions. Bottom, boxplots indicate the median, 25th and 75th percentile (box edges), and 1.5 times the interquartile range (whiskers) of the two distributions.

Inhalations periodically activated most PCx neurons in a sequential manner, and faster sniffs appeared to enhance the activity of some neurons while decreasing that of others (Figure 1D); as a result, PCx activity tracked the phase and speed of each inhalation (Figure 1E-F and Figure S1C-D). 29.6% of the recorded neurons reliably responded to at least one odor concentration (195 of 660 neurons; 6 mice). How fast an animal inhaled influenced the response amplitude during inhalation in 71.3% of those neurons (139 of 195 neurons; Figure 1G-I). This proportion was a lower-bound estimate based on the responses to a panel of only five odors and two concentrations. Remarkably, the impact of a faster inhalation was not uniform across odor responses: 33.2% of the odor responses increased with a rapid inhalation, whereas 22.7% decreased in amplitude. This sniff-driven variability in odor responses was already significant during the first 70 ms of an inhalation (Figure S1E). Because GABAergic neurons providing feedback inhibition tend to exhibit narrower action potentials and faster activity^23^, we used these features to tentatively distinguish them from the other excitatory and feedforward inhibitory neurons. After sorting neurons in regular spiking (RS, putatively excitatory or feedforward inhibitory) and fast spiking (FS, putatively feedback inhibitory), we observed similar effects of the inhalation speed in both types (Figure S1F).

On average, sniffs caused a response amplitude change bigger than that caused by a ten-fold increase in odor concentration in the odor-responsive neurons (Figure 1J). Thus, changing the inhalation speed generated a large amount of response variability that could impair the discriminability of odor concentrations on an inhalation-by-inhalation basis. Figure 1K illustrates this problem in an example PCx neuron. The raster plot and the peri-event time histogram (PETH) in the left column show the responses to two concentrations of the same odor during slow breathing. This neuron generated more action potentials when the odor was delivered at the highest of the two concentrations. The right column shows the responses to the lowest concentration upon a slow or a fast inhalation. The response amplitude to the low concentration during a rapid inhalation was indistinguishable from that measured during a slow inhalation of the more concentrated odor. To assess how varying the inhalation speed affected concentration encoding across all odor-responsive neurons, we calculated a concentration discriminability index for each neuron (CDI; see *Methods*). The CDI was computed using the odor responses during slow, fast, or both types of inhalation. Mixing responses from slow and fast inhalations decreased the CDI, consistent with a more extensive response variability than within the same inhalation speed (Figure 1L).

### PCx neurons randomly mix olfactory and non-olfactory inputs during inhalations

Sniffing could alter odor responses by changing the odorant deposition onto the olfactory epithelium. However, we found odor responses that increased in amplitude with the concentration but decreased with the inhalation speed and others with the opposite pattern (Figure 2A). These observations suggested that odor responses also vary with the inhalation speed because of a non-olfactory input. To assess the relationship between this non-olfactory input and concentration encoding, we fitted a generalized linear model (GLM) to the odor responses of PCx neurons recorded upon different odor concentrations and inhalation speeds. The GLM included three factors: odor concentration (0%, 0.01%, 0.1%, and 1% v/v), inhalation speed (slow or fast), and the interaction between inhalation speed and concentration (Figure 2B). The inhalation speed factor can be interpreted as a non-olfactory input that is combined with the odor input. One way to interpret the multiplicative interaction between inhalation speed and odor concentration is the potential effect of the airflow rate on the concentration profile of the odorant in the nasal cavity^4^. Other types of interaction cannot be excluded, however. For example, sniffs could be associated with attentional mechanisms that change the gain of the odor response; the interaction weight should capture this effect. To identify the factors that were more relevant for the odor responses of a neuron, we regularized the GLM with an elastic net method. This model explained 91.4% of the variance in the mean number of spikes generated during the first 180 ms of odor inhalation across all neurons.

**Figure 2.**
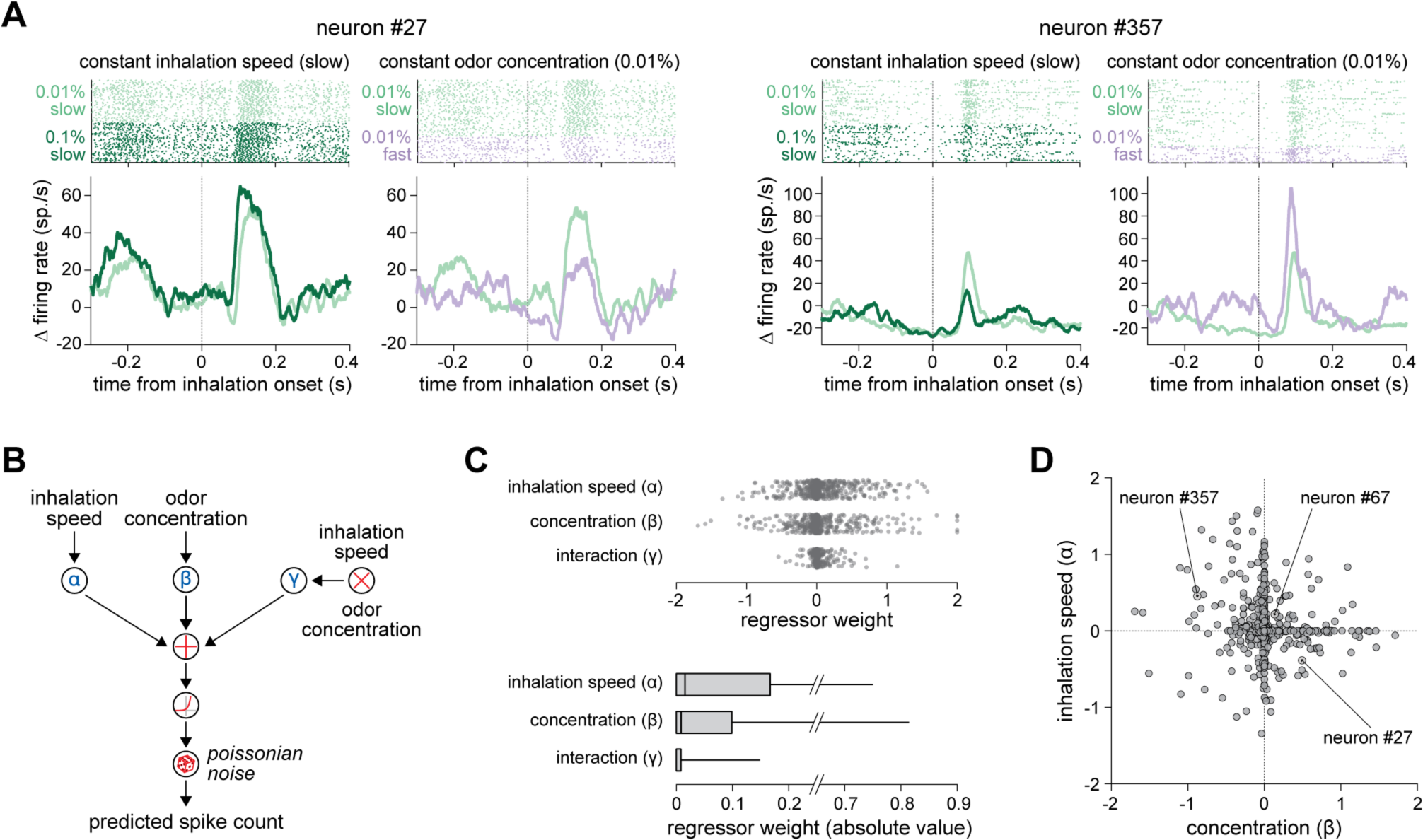
PCx neurons randomly mix olfactory and non-olfactory inputs during inhalations. **(A)** Examples of responses from two neurons with independent sensitivities to odor concentration and inhalation speed. For each neuron: left, raster plot and PETHs of the responses to 0.01 % (light green) and 0.1 % (dark green) v./v. averaged across all inhalations; right, raster plot and PETHs of the responses to slow (light green) and fast (light purple) inhalations of 0.01% v./v. odor. **(B)** Schematic of the generalized linear model (GLM) used to predict the odor response based on inhalation speed (slow and fast), odor concentration (0, 0.01, 0.1, 1% v./v.), and their interaction. The GLM was regularized with an elastic-net method. **(C)** Top, distribution of the regularized regressor weights for inhalation speed, odor concentration, and their interaction (n = 928 neuron-odor pairs; 2 odors; 464 neurons; 4 mice). Bottom, boxplot of the absolute values of the regularized regressor weights indicating the median, 25th and 75th percentile (box edges), and the 95^th^ percentile (whisker). **(D)** Scatterplot of the concentration and inhalation speed regressor weights. Each data point is a neuron-odor pair (n = 928 neuron-odor pairs; 2 odors; 464 neurons; 4 mice). Neurons #27 and 357# of Figure 2A and neuron #67 of Figure 1K are indicated.

The GLM analysis revealed that the interaction between inhalation speed and concentration response had a small effect on the odor response amplitude during an inhalation (Figure 2C). Instead, the non-olfactory input had a more significant additive or subtractive effect (Figure 2C). Importantly, the sign of the weight for the non-olfactory input and the odor concentration factors could differ in the same neuron (Figure 2D). The non-olfactory input caused by a fast inhalation could increase or decrease the odor response amplitude regardless of how the neuron responded to a higher concentration of that odor. Consequently, the combination of non-olfactory and olfactory sensitivities was heterogeneous across PCx neurons (Figure 2D). We found a similar distribution of non-olfactory sensitivities in RS and FS neurons (Figure S2A-B). Moreover, restricting the analysis to the early 70 ms of inhalation also gave similar results (Figure S2C). Thus, the non-olfactory input appeared to be the leading cause of the odor response variability observed at different inhalation speeds.

### A mechanosensory input signals the nasal airflow rate to the PCx

Sniffing and blowing non-odorized air into the nostrils have been shown to activate the PCx^16^. Therefore, the inhalation speed may affect the odor responses in PCx through a mechanosensory input originating in the nasal cavity. Interestingly, however, the results of the GLM analysis suggested that a sniff could increase the odor response of some neurons but also decrease it in others. To further investigate this observation, we directly assessed how the air passage through the nose affects the spiking activity of PCx neurons.

We first performed a tracheostomy in anesthetized mice to eliminate air passage through the nose during respiration (Figure 3A). This procedure interrupted the normal entrainment of PCx activity to the breathing rhythm as expected (Figure S3A). Then, we mechanically stimulated the nasal epithelium with 150-ms-long airflow pulses of odorless air delivered a 1 Hz. We tested five different airflow rates within the range of an awake mouse inhalation^11,19,30,31^. 57.1% of neurons responded to the artificial mechanosensory stimulation of the nasal cavity. The response latencies varied across neurons, like in normally breathing mice upon inhalation. The response amplitude varied with the airflow rate. Some neurons increased their response amplitude as the flow rate increased, whereas others fired fewer action potentials (Figure 3B-C). Faster airflow also decreased the response latency without significantly changing the sequential activation order across neurons (Figure 3B and Figure S3B).

**Figure 3.**
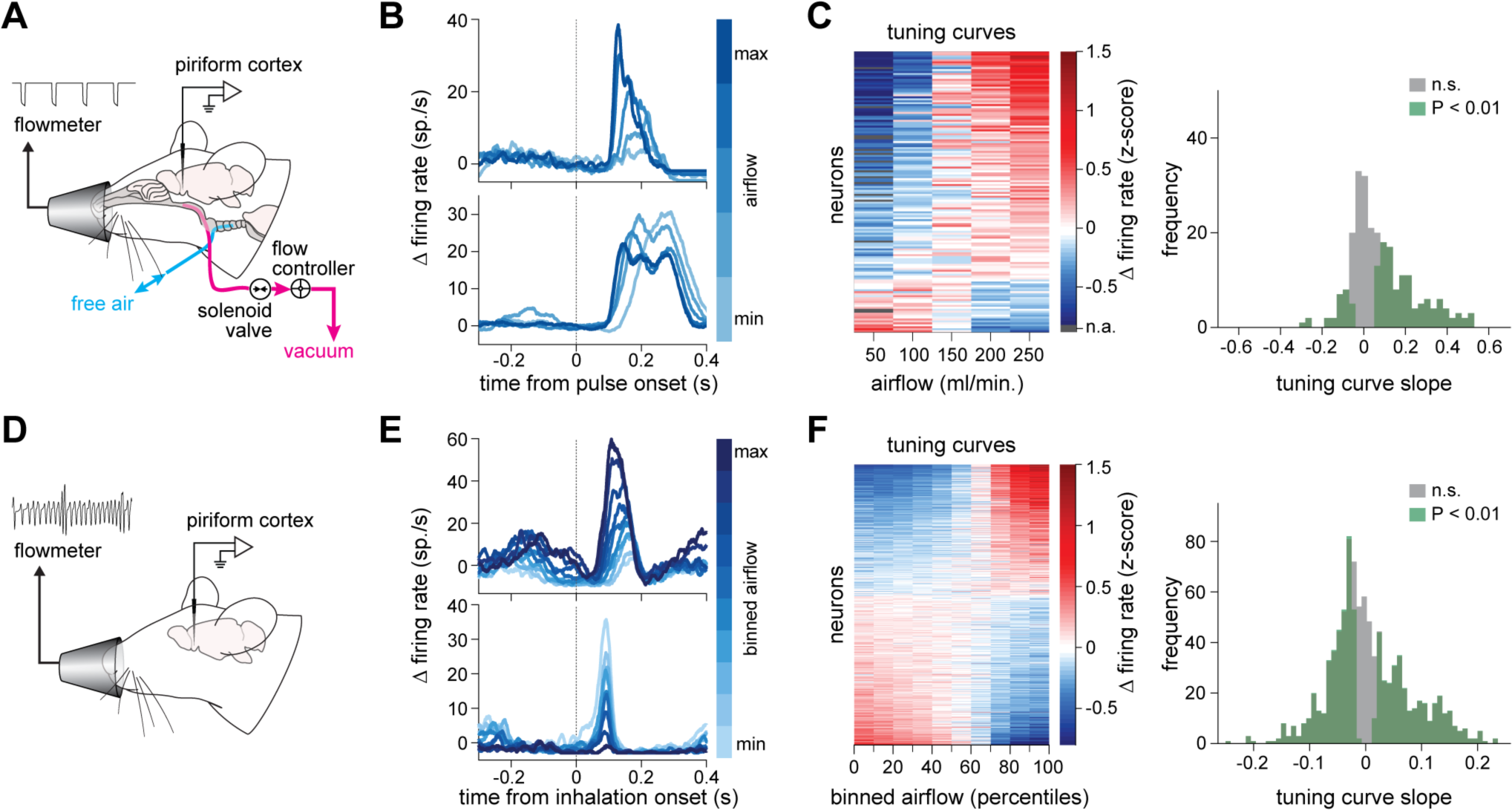
A mechanosensory input signals the nasal airflow rate to the PCx. **(A)** Experimental setup for artificial airflow stimulation of the nasal cavity in anesthetized mice. **(B)** PETHs of two example neurons responding to five different airflow velocities. **(C)** Left, raster map of airflow velocity tuning curves (n = 144 velocity-tuned neurons of 252; 3 mice). Right, distribution of the slopes of the airflow tuning curves. Shaded area, slopes significantly different from zero. Positive slope: 117 neurons; negative slope: 27 neurons (*P* - value < 0.01, t-test). **(D)** Experimental setup for PCx recordings paired with spontaneous breathing monitoring in awake mice in the absence of odors. **(E)** PETHs of two example neurons responding to ten different peak airflow speeds. **(F)** Left, raster map of the tuning curves for the peak airflow velocity (n = 748 velocity-tuned neurons of 955; 10 mice). Right, distribution of the tuning curve slopes like in (C). Positive slope: 350 neurons; negative slope, 398 neurons (*P*-value < 0.01, t-test).

That these mechanosensory responses were not an artifact of the artificial stimulation of the nasal cavity in an anesthetized mouse is suggested by the finding of similar mechanosensory tuning curves in awake and normally breathing mice (Figure 3D). For each mouse, we divided the range of the peak airflow rates measured during all inhalations into ten bins. All bins had an equal number of inhalations. Then, we measured the mean amplitude of the spiking activity during the inhalations within each bin. This analysis revealed that 78.3% of the PCx neurons (77.2% of the odor-responsive neurons) encoded the peak inhalation airflow rate with a monotonic change in spiking activity after the inhalation (Figure 3E). Sorting the airflow tuning curves of all neurons based on their slopes revealed a push-pull organization of the inhalation speed responses again; that is, 36.7% of the neurons increased the amplitude of their responses when the inhalation speed increased, and another 41.7% decreased their responses (Figure 3F).

What was the source of the mechanosensory input? OSNs are excited by pressure changes, and blocking OSNs’ responses disrupts the typical synchronization between respiration and the local field potential recorded in the OB^20^. Thus, OSNs are considered a source of mechanosensory information for the olfactory system. Nonetheless, our results could not exclude the possibility that trigeminal sensation could also provide nasal airflow information to the PCx. For example, the trigeminal input may afford the sensitivity to the airflow rate, and the OSN input may only give a bulk activation indicating the inhalation onset. Intriguingly, a branch of the trigeminal nerve called the anterior ethmoidal nerve (AEN) innervates the olfactory epithelium and the OB as well^32^. The fibers of the AEN respond to chemical and mechanical stimulation^33^. Therefore, the AEN afferences could sense the airflow rate inside the nasal cavity and broadcast this information to the PCx through the OBs. To test this possibility, we sectioned the AEN (Figure S3C). In a subset of mice, we occluded the contralateral nostril to prevent information leakage from the contralateral side. Severing the AEN left intact the responses of PCx neurons to inhalations. The tuning curves for the airflow rate after the neurectomy remained similar to those of control mice (Figure S3D-E). These data indicate that PCx neurons encode the inhalation airflow rate in the nasal cavity through a mechanosensory input that most likely arises from the OSNs.

### A top-down input supplies behavioral state information during a sniff

While the above results show that the PCx encodes the airflow rate inside the nasal cavity, they do not eliminate the possibility that top-down inputs could also account for the activity changes caused by faster sniffing in PCx. Sniffing characterizes active exploration and arousal states^29,34^. Therefore, top-down and neuromodulatory inputs associated with those behavioral states may affect the activity of PCx neurons during a sniff. The responses to these inputs could be hard to distinguish in normal conditions because they would likely be mixed with the mechanosensory responses. Moreover, although our artificial ventilation experiments uncoupled the respiratory activity from the mechanical stimulation of the nasal cavity, they also prevented voluntary sniffing because they were performed under anesthesia. To isolate the possible effect of top-down inputs during a sniff, we removed both OBs in a group of mice and recorded the activity of PCx neurons during wakefulness. We examined the responses to the first fast sniff after a sequence of at least five slow inhalations and to a regular slow inhalation in the absence of odors.

In bulbectomized mice, PCx neurons did not respond to either odors (Figure S4A) or slow inhalations (Figure 4A). These results confirmed that the bulbectomy blocked peripheral inputs to the cortex. Nonetheless, 17.2% of PCx neurons still responded to sniffs. For comparison, this proportion was 29.7% in control mice. However, the sniff responses differed from those of control mice (Figure 4B-E). In bulbectomized mice, the response amplitude was smaller and less reliable across sniffs than in control mice. Consistently, a linear classifier reading the PCx activity pattern during an inhalation could distinguish a sniff from a regular inhalation in both control and bulbectomized mice; however, the decoding accuracy after bulbectomy was significantly lower than in control mice (Figure 4F). Remarkably, the sniff response revealed by the bulbectomy seemed to start rising several tens of milliseconds before the nominal onset of the sniff. Since responses to slow inhalations were absent in the bulbectomized mice, this early response could not be attributed to the previous inhalation. Moreover, we observed similar sniff responses in the motor cortex of mice with intact OBs (Figure S4B-D). Together, these data demonstrate that sniffs can be preceded by a global top-down signal that may further affect odor responses in PCx.

**Figure 4.**
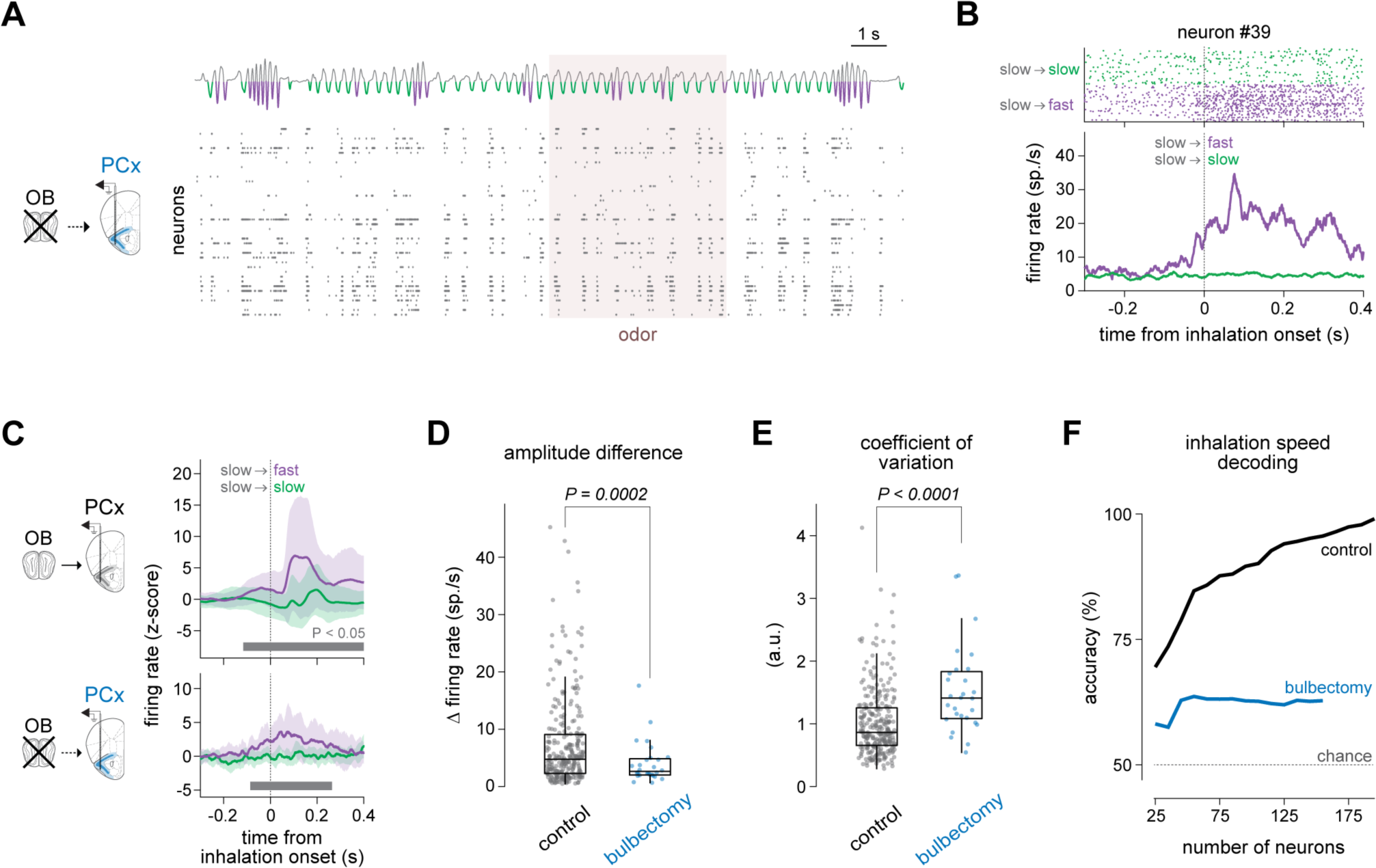
A top-down input supplies behavioral state information during a sniff. **(A)** Example of simultaneous PCx activity and respiratory airflow speed in an awake mouse after a bulbectomy. External flow sensor signal (top) and spiking raster plot (bottom). Shaded area, odor period (5 seconds). Slow and fast inhalations are color-coded in green and purple, respectively. **(B)** Spiking activity during slow and fast inhalations in an example neuron after a bulbectomy. Raster plot (top) and PETH (bottom) are shown. The responses to the first sniff after at least five slow inhalations and to a slow inhalation are color-coded in purple and green, respectively. **(C)** Average PETHs of neurons preferring a sniff over a slow inhalation in mice with intact OBs (top; 284 of 955 neurons; 10 mice) and in bulbectomized mice (bottom; 28 of 162 neurons; 4 mice). Shaded area, mean ± s.e.m. The bar below the PETH indicates when the sniff responses are significantly bigger than the responses to a regular inhalation (Benjamini-Hochberg adjusted *P*-value < 0.05, one-sided t-test). The average responses to the first sniff after at least five slow inhalations and to a slow inhalation are color-coded in purple and green, respectively. **(D)** Difference between the amplitudes of the responses to the first sniff and a slow inhalation in control and bulbectomized mice (*P*-value = 0.0002; two-sided t-test with unequal variances). Boxplots indicate the median, 25th and 75th percentile (box edges), and 1.5 times the interquartile range (whiskers). **(E)** Coefficient of variation of the response amplitude across first sniffs in control and bulbectomized mice (*P*-value < 0.0001; two-sided t-test). Boxplots indicate the median, 25th and 75th percentile (box edges), and 1.5 times the interquartile range (whiskers). **(F)** Decoding accuracy of inhalation speed as a function of the neurons included in the analysis; classifiers used the activity of a pseudo-population of PCx neurons in the control condition (black line) and after bulbectomy (blue line).

### Odor and inhalation information are independent

An important result of the GLM analysis was that odor and inhalation speed sensitivities appeared randomly distributed among odor-responsive PCx neurons. This heterogeneity in sensitivity raises the possibility that the representations of the two modalities do not interfere at the population level (see *Methods* for a formal derivation in linear encoding systems). PCx represents the identity and concentration of a given odor with a specific pattern of activation of ensembles of neurons or, equivalently, with population vectors^24,25,28,35,36^. Figure 5A shows the PCA embedding of the population response vectors to three concentrations (0.01, 0.1, 1% v/v) of an odor upon slow and fast inhalations (see also Figure S5A for the inhalation-by-inhalation population responses in individual mice and Figure S5B for a different odor). Increases in concentration result in scaling the population vector magnitude without changing vector direction, as already observed in prior work^22,23^. Since the sniff response of an odor-responsive neuron weakly correlated with that of other odor-responsive neurons, the direction of the population vector response to an increase in inhalation speed did not overlap with the direction of the concentration population response. In other words, inhalation speed information did not affect the encoding of the odor concentration.

**Figure 5.**
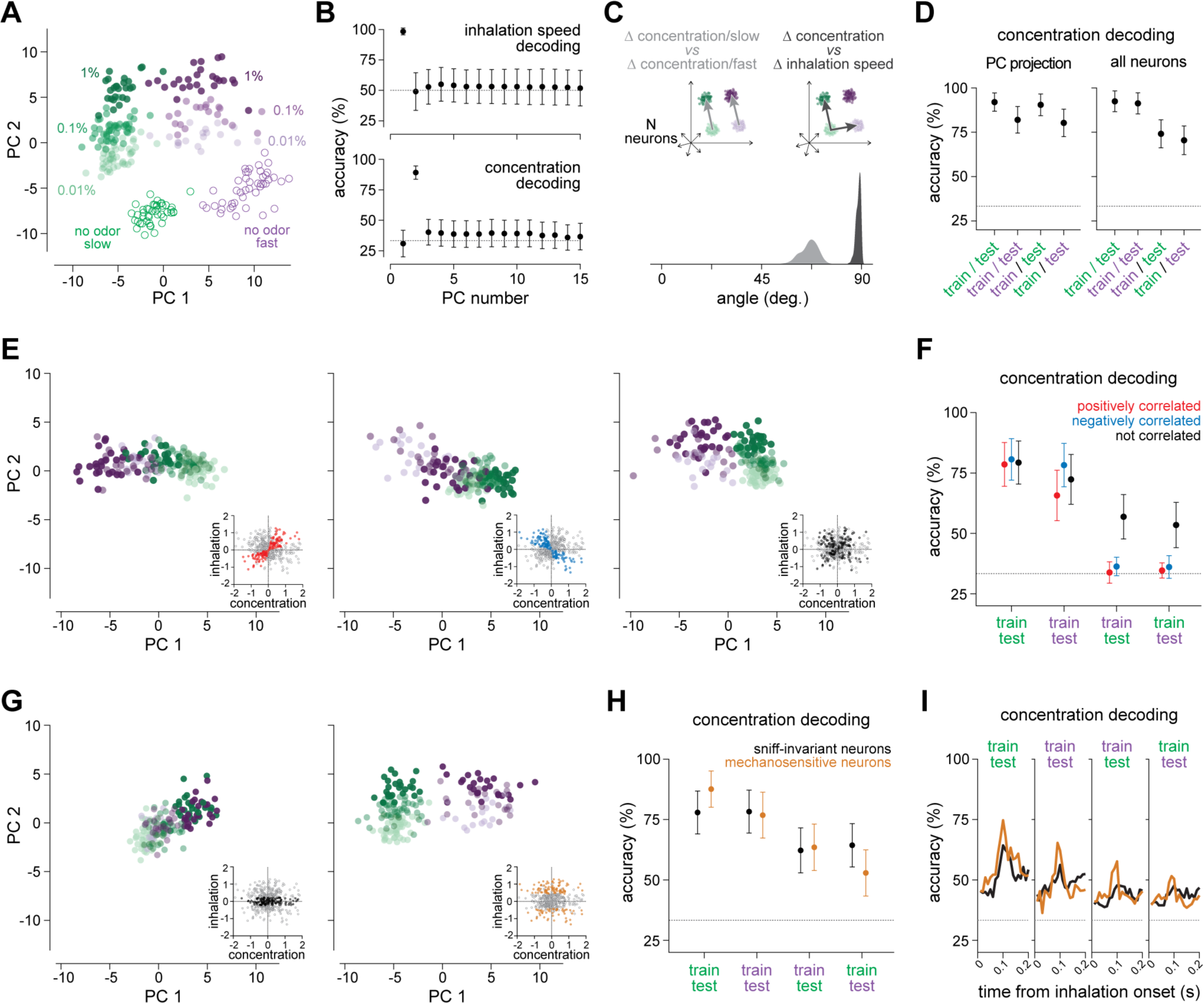
Odor and inhalation information are independent. **(A)** PCA embedding of pseudo-population responses (464 neurons, four mice) to three different concentrations of an example odor during slow (green) and fast (purple) inhalations. Each dot is a response in a 180 ms window starting at inhalation onset. Empty dots represent inhalation responses in the absence of odors. A random sample of all responses is shown. **(B)** Decoding accuracy of inhalation speed (top) and odor concentration (bottom) using the projection of the pseudo-population odor responses in (A) on each of the first fifteen PCs. Mean ± standard deviation is shown. **(C)** Angle between vector pairs representing concentration changes (light grey) and vector pairs representing a concentration change and an inhalation speed change (dark grey). **(D)** Left, concentration decoding accuracy of *cis*- and *trans*-decoders using the pseudo-population response projections onto the second and third PCs. Right, concentration decoding accuracy of *cis*- and *trans*-decoders using non-reduced population responses. *Cis*-decoders were trained on slow (green) or fast (purple) inhalations and tested on held-out slow or fast inhalations, respectively. *Trans*-decoders were trained on slow (green) or fast (purple) inhalations and tested on held-out fast or slow inhalations, respectively. Mean ± standard deviation is shown. **(E)** PCA embedding of concentration responses for three pseudo-populations with positive (left), negative (center), and no correlation (right) between inhalation and concentration sensitivity. Insets: scatterplots of inhalation and concentration regression coefficients for all the neurons of the same pseudopopulation used in (A). Red, neurons with positive correlation; blue, neurons with negative correlation; black, neurons without correlation. An equal number of neurons was randomly sampled from each sub-population (n = 143 neurons). A random sample of all responses is shown. **(F)** Concentration decoding accuracy of *cis*- and *trans*-decoders using the three neural sub-populations in (E). Mean ± standard deviation is shown. **(G)** PCA embeddings of concentration responses using a pseudo-population with (left) and without (right) sniff-invariant neurons. Mechanosensory sensitivity was estimated as the inhalation regression coefficient of a two-factor (inhalation speed and concentration) linear model fitted to single-neuron responses. Insets: scatterplots of the inhalation and concentration regression coefficients for all the neurons of the same pseudopopulation used in (A). The subset of neurons in the population response vector is highlighted by color: black, sniff-invariant neurons; orange, mechano-sensitive neurons. An equal number of neurons was randomly sampled from each sub-population (n = 143 neurons). A random sample of all responses is shown. **(H)** Concentration decoding accuracy of *cis*- and *trans*-decoders inhalation speed using sniff-invariant (black) and mechano-sensitive (orange) neurons. Mean ± standard deviation is shown. **(I)** Concentration decoding accuracy within and across inhalation speed over the time course of an inhalation (*P*-value > 0.05; two-way ANOVA). An equal number of neurons was randomly sampled from each sub-population (n = 143 neurons).

To confirm this qualitative observation, we used an linear support vector machine (lSVM) to decode either the inhalation speed or the odor concentration from the projections of the PCx responses on the first 15 PCs (Figure 5B and Figure S5C-D). Linear decoders could discriminate the inhalation speed with high accuracy using the projection of neural activity onto the first PC. In contrast, decoders could not classify an odor concentration using only the activity explained by the first PC. Moreover, decoders could not classify an odor identity using only the activity explained by the first PC (Figure S5E), further suggesting that the variance explained by the first PC does not encode olfactory information. Conversely, linear classifiers decoded the odor concentration and identity with high accuracy when they used neural activity along the second and third PCs (Figure 5B, and Figure S5C,E). However, classifiers could not discriminate the inhalation speed using the second and subsequent PCs.

Importantly, we expected that how PCx encoded odor concentration would not depend on inhalation speed. To test this hypothesis, we used linear discriminant analysis to find two directions in the population neural activity that best tell apart odor concentrations during slow or fast breathing. We found that, even though these two directions were different (as expected, given the high dimensionality of the population response), they were more similar than the directions that tell apart how fast a mouse breathes in from how concentrated the odor is (Figure 5C).

### The distributed representation of odor concentration is sniff-invariant

These results suggested that the cortical representation of odor concentration could be sniff-invariant. We thus trained an lSVM to decode an odor concentration using a subset of slow or fast inhalations; then, we tested the performance of the same decoder using held-out inhalations with the same (*cis*-) or other speed (*trans*-decoding). A decoder using PCx response projections on the second and third PCs exhibited similar *cis*- and *trans*-decoding performances, suggesting that PCx has sniff-invariant representations of odor concentration (Figure 5D and Figure S5F-G). Next, we tested the linear decoders on the untransformed PCx responses, thus removing the possibility that the decoder could leverage the denoised representations provided by PCA; this *trans*-decoder could still predict the odor concentration, generalizing across inhalation speeds (Figure 5D and Figure S5F). We observed similar results when the decoders used spike counts summed over 70 or 180 ms windows and spike counts for concatenated 10 ms or 10° bins over a respiratory cycle (Figure S5G-H).

How well the concentration information was preserved across inhalation speeds was determined by how heterogenous the combination of inhalation speed and olfactory tuning was across neurons. To confirm this, we performed PCA and decoding analysis on neural sub-populations with correlated sensitivity to non-olfactory and olfactory inputs, thus removing the heterogeneous distribution of non-olfactory and olfactory responses. PCA embeddings of the odor responses showed that representations of odor concentrations become ambiguous across inhalation speeds in these sub-populations (Figure 5E). Thus, a *trans*-decoder performed at the chance level if it could only use the activity of a neural population with correlated sensitivity for non-olfactory and olfactory inputs (Figure 5F). Instead, using a population with uncorrelated selectivity disentangled the representations of inhalation speed and odor concentration, thus allowing a *trans*-decoder to distinguish odor concentrations across inhalation speeds (Figure 5E-F).

Our dataset also included neurons that did not change their odor responses with the inhalation speed. A downstream reader could privilege the information from these sniff-invariant neurons to obtain a stable readout of the odor concentration across inhalation speeds. We determined whether a population of neurons insensitive to the inhalation speed could afford more sniff-invariant information than the rest of the neural population. To select the sniff-invariant neurons, we assessed the sensitivity to the inhalation speed of each neuron in our dataset. To this aim, we fitted the inhalation-by-inhalation odor responses with a linear regression model with a concentration and an inhalation speed regressor. Neurons with an inhalation speed regressor coefficient within 0 ± 0.75 standard deviations were included in the sniff-invariant population. In contrast, an equal number of neurons with an inhalation speed regressor coefficient beyond ± 1.5 standard deviations from zero were included in a population that we denoted as mechano-sensitive (Figure 5G). The two populations of sniff-insensitive and mechano-sensitive neurons had similar distributions of concentration regressor coefficients (Figure S5I). We compared the classification accuracies of linear *trans*-decoders using sniff-invariant and mechano-sensitive populations. This analysis revealed that a population of mechano-sensitive neurons carries at least as much sniff-invariant information as a population of only sniff-insensitive neurons (Figure 5H).

We also examined the possibility that sniff-insensitive neurons could provide more sniff-invariant information than mechano-sensitive neurons during the earliest part of an inhalation. To this aim, we employed the *cis*- and *trans*-decoding approach in twenty consecutive time bins (10 ms/bin), tiling the first 200 ms of an inhalation. Linear decoders using only mechano-sensitive or sniff-insensitive neurons performed similarly throughout this inhalation window (Figure 5I). Together, these results confirm that sniff-invariant concentration information is distributed across the activity pattern of the odor-responsive PCx neural population.

### Airflow information affords tolerance to flow-dependent concentration fluctuations

Our data indicated that adding a mechanosensory response to olfactory responses does not significantly alter the representation of odor concentration in PCx. In fact, adding the inhalation speed information teases apart the representations of the odor concentration under different inhalation speeds. Therefore, this coding strategy may allow a linear decoder to distinguish an environmental increase in concentration from one induced by a nasal airflow change. This way, the decoder can learn that the two activity patterns evoked by an odor upon slow and fast inhalations represent, in fact, the same concentration (Figure 6A). To test the robustness of this coding strategy, we simulated the responses of a population of neurons with varying levels of inhalation speed-dependent concentration change and mechanosensory input. To generate the surrogate distribution of responses, we used the odor and inhalation covariate weights obtained by fitting our recorded neural responses with a linear regression model. We simulated different levels of interaction between airflow and odorant deposition by assuming that the inhalation-driven change in concentration (ΔIC) is proportional to the initial concentration and the inhalation speed, as per prior fluid dynamics modeling^4^ (see *Methods*). We varied ΔIC over a two-fold range to approximate the natural nasal airflow rate range^37^. PCA embedding showed that odor representations were ambiguous in this simulated population when there was no inhalation input (Figure 6B); for example, an increase in odor concentration could not be distinguished from an increase in inhalation speed. Consistently, decoders trained on the responses during slow and fast inhalations classified the odor concentration worse than decoders trained using only one inhalation speed as the simulated influence of the airflow on the concentration increased (Figure 6C). However, supplementing this population with mechanosensory responses was sufficient to reduce ambiguity and retrieve the actual concentration for any ΔIC (Figure 6A-C).

**Figure 6.**
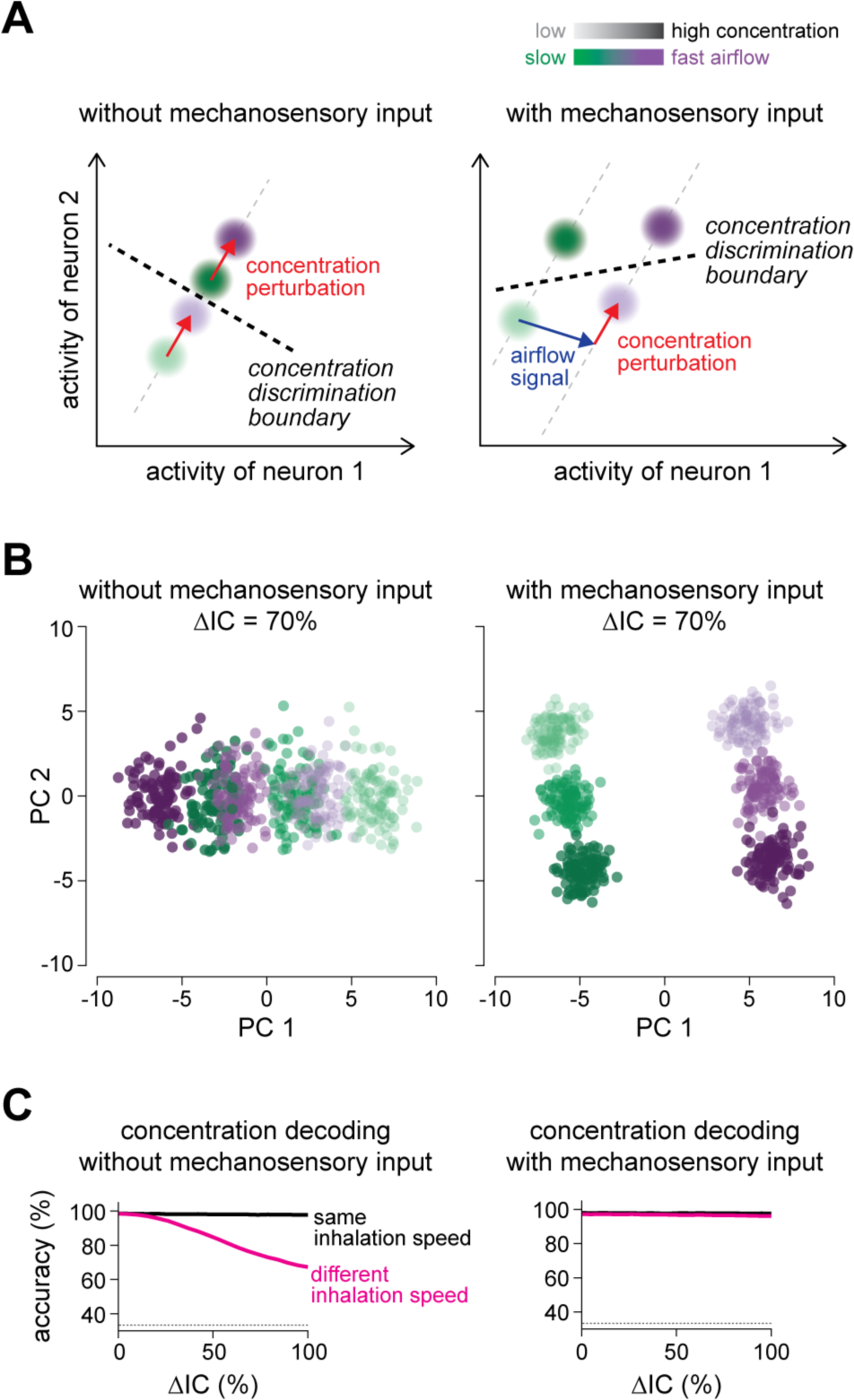
Airflow information affords tolerance to flow-dependent concentration fluctuations. **(A)** Example of how adding an independent mechanosensory signal distinguishes the inhalation-induced concentration fluctuation from an inhalation-independent change in concentration. The X- and Y-axis represent the activity of two neurons. The dots represent the inhalation-by-inhalation population responses to an odor. The saturation indicates the odor concentration. The hue indicates the inhalation speed. A faster inhalation carries more odorant molecules per unit of time, increasing the concentration in the naris. Left, the representations of a higher concentration caused by a sniff (light purple) or an external event during a regular inhalation (dark green) are indistinguishable. Right, adding an independent airflow signal opens a new subspace to represent odor concentrations during a sniff. Thereby, a linear decoder can learn a concentration discrimination boundary tolerant to the concentration fluctuations caused by the sniff. **(B)** Inferred effect of the mechanosensory input on the concentration representation in the presence of an inhalation-driven alteration of the odor concentration. Left: PCA embedding of population concentration responses with a 70% inhalation-driven concentration change (ΔIC) and without inhalation input. Right: PCA embedding of population concentration responses with a 70% ΔIC and mechanosensory input. Population responses were simulated using the inhalation and concentration regression coefficients estimated from the actual data to generate surrogate single-neuron responses. The ΔIC was artificially added to the concentration covariate of the model as a term proportional to the actual concentration and the inhalation speed. **(C)** Concentration decoding accuracy for increasing levels of inhalation-dependent ΔIC without (left) and with (right) mechanosensory input within (black) or across (magenta) different inhalation speeds.

## DISCUSSION

Sniffing alters odor responses at the early stages of olfactory processing^4,5,44,17,31,38–43^. These observations question whether the olfactory system can afford a stable representation of odor concentration despite variability in inhalation speed. Our study shows that concentration information is resistant to changes in inhalation speed. The explanation is that most PCx neurons mainly respond to a mechanosensory input that indicates the nasal airflow rate, sums with the odor input but is not necessarily correlated with the latter. Thus, the airflow and concentration sensitivities combinations are heterogeneous among the PCx neurons activated by a given odor. At the population level, this heterogeneity has two advantages. First, the mechanosensory response does not interfere with concentration encoding. Second, the mechanosensory response differentiates the representation of a potential concentration increase caused by a quick breath from that of an environmental rise. Thus, a linear decoder reading the population activity can discriminate the true odor concentration using the same code for both slow and fast inhalations.

Our artificial inhalation experiments support the hypothesis that the PCx encodes information about inhalation dynamics through reafferent mechanosensory stimuli from the olfactory epithelium. Furthermore, interrupting the information flow through the OBs deprived the PCx of most inhalation information. That PCx neurons respond to inhalations is not unusual^16,18,45,46^. However, our findings uncover a previously unappreciated representation of the nasal airflow speed in the PCx. The mechanosensory tuning curves that we measured showed a push-pull arrangement across the population of PCx neurons, meaning that a faster inhalation increased the response of some neurons while decreasing that of others. Such a distribution of tuning curves is known to stabilize the average firing rate and keep the information rate constant across the full range of nasal airflow velocities experienced by the mouse^47^. We propose that such a bidirectional organization of the mechanosensory responses is important because, combined with a random sampling of odor and mechanosensory inputs from the OBs, it expands the possible combinations of sensitivities to the odor and inhalation inputs, warranting a more robust tolerance to varying inhalation speeds in PCx. If downstream brain areas also use this rich respiratory information for other goals remains to be investigated.

The observation that the inhalation response amplitude could decrease with the inhalation speed was unexpected. Based on previous work^20^, increasing the flow rate would be predicted to increase the mechanosensory responses of OSNs. Blowing air into the nostrils was shown to activate the PCx^16^. Furthermore, faster sniffs reduce the response latency of MT neurons to an inhalation^4,5^; because PCx neurons are thought to transform an increase in MT inputs synchrony into a rise in firing activity^48^, we expected that a faster sniff would increase the response of PCx neurons. Instead, we found that quick sniffs reduced the response amplitude. Notably, we observed response amplitude decrements even after the first fast sniff of a sniffing bout, which rules out the possibility of adaption to repeated sniffs. Thus, our finding suggests the involvement of an inhibitory circuitry that could be specifically sensitive to sniffing. Where is this circuitry? A fast inhalation may enhance some mechanosensory inputs and weaken others already at the level of the olfactory glomeruli. Indeed, sniffing can also reduce the activity of mitral cells^5^. Quick breaths may trigger an asymmetric interglomerular inhibition or pre-synaptic inhibition within the same OSN channel, similar to mechanisms previously described for odor processing^49–51^. However, we cannot exclude that sniffing dampens the strength of mechanosensory responses in some PCx neurons through cortical feedforward and feedback inhibitory circuits^23,46,52–55^.

Removing the OB reduced the information about the inhalation speed in PCx. However, some residual information was still present, suggesting that top-down inputs may modulate PCx activity during sniffing. Indeed, the PCx receives projections from other forebrain areas, such as the orbitofrontal cortex^56^, the amygdala^57^, and the parahippocampal region^58,59^. Furthermore, cholinergic inputs increase the excitability of PCx neurons^60^, and the stimulation of the locus coeruleus increases the responses of PCx neurons to inhalations^61^. These top-down inputs may influence odor processing during active behavior^27,29^. For example, in mice performing an olfactory task, the activity of PCx neurons is sensitive to the different epochs of the task as well as the odors^62,63^. Nevertheless, the computational role of top-down inputs to PCx is unknown. The responses to fast inhalations after the bulbectomy may represent the widespread effect of an arousal state^29^. Yet, since we only monitored the respiratory activity, we cannot exclude that a small group of PCx neurons encoded other types of movement, as observed in other sensory cortices^64–72^. Interestingly, we noticed that, in bulbectomized mice, the sniff response started before the inhalation onset. Moreover, this response was less reliable than the mechanosensory response across rapid inhalations. A possible explanation for such variability is that it reflected the variance between fast inhalations. Some of this inhalation speed variability may depend on whether the faster inhalation was caused by an autonomic change in respiratory activity or an explorative sniff^29^. These considerations raise the possibility that the top-down input we observed may represent a global preparatory signal working in concert with the mechanosensory feedback from the nasal cavity only during voluntary sampling.

The information provided by the airflow signal differentiates concentration representations upon different inhalation speeds. However, it cannot correct the concentration change caused by varying the inhalation speed in and by itself. Although the mechanosensory feedback is the same regardless of which odor an animal smells, the concentration change caused by a change in inhalation speed is likely different for different scents. Indeed, how much odorant crosses the olfactory mucosa depends on the airflow velocity and the odorant’s air-water partition coefficient^73,74^. Thus, the correspondence between two representations of the same environmental concentration upon two inhalation speeds has to be learned from experience. In most instances, the concentration of an odorous source (for example, the octanal molecules in an orange peel) tends to be stable during an olfactory experience. We hypothesize that the brain could learn to generalize across inhalation speeds by taking advantage of the temporal contiguity of an odorant source during consecutive inhalations. A similar, unsupervised learning strategy might work even in a more naturalistic scenario involving variable odor plumes and odorant mixtures. Further work will be required to investigate how the olfactory circuits exploit the spatiotemporal statistics of the olfactory environment and fully understand the relationship between experience and tolerance to self-generated stimulus variability.

Our work reveals a novel solution to the sensory disturbances caused by movements. This solution does not rely on inhibitory corollary discharges, but leverages the computational benefits of population codes. Such a coding strategy is so general that it is likely to be used by other sensory systems. Indeed, the co-existence of independent motion and sensory signals is not a unique feature of the olfactory system. Orthogonal dimensions for encoding movements and sensory stimuli have been found in other sensory cortices, such as the primary visual cortex (V1)^66,71,72^. However, the function of these movement signals is a matter of debate^64,65^. In recent work, it was observed that V1 neurons independently encode the direction of a drifting visual grating and a saccade. It was proposed that this encoding strategy may “scramble” the image representation during the saccade, thus suppressing the perception of the retinal image drift caused by the saccade^70^. Here, we observe that the motion-related signal does not destroy the olfactory information in PCx. Rather, the sniff signal lawfully reconfigures the concentration representations according to the nasal airflow. Thus, a downstream area can learn an optimal decoding scheme that retrieves the true concentration, like with a conversion chart (see the model in Figure 6A). The motion signals in V1 could have a similar function. Interestingly, psychophysical experiments in humans suggest that reafferent mechanosensory signals reduce the perceived extent of the visual smear generated by passive head and eye movements^75^; notably, these observations also offer a functional interpretation of the additive inputs that the mouse V1 receives from the vestibular system^68,69,76^. Mechanosensation is the analysis of movement; thus, embedding independent mechanosensory reafferences into distributed sensory codes could be a canonical neural design to parse and reject the consequences of sensory organ movements on sensation.

## AKNOWLEDGMENTS

We thank S.R. Datta, J.A. Assad, T. Fellin and G. Iannetti for comments on this work. We thank the animal facility staff for assistance with mice husbandry. We thank M. Libera for technical assistance. We thank R. Poli, P. Battistoni and S. Maestrelli for administrative support. G.I. holds a Giovanni Armenise-Harvard Foundation Career Development Award; L.P. is supported by an EMBO Postdoctoral Fellowship.

## AUTHOR CONTRIBUTIONS

Conceptualization: G.I.

Methodology: F.M., P.P., A.A.D., G.I.

Software: A.A.D., L.P., P.P., G.I.

Formal analysis: A.A.D., L.P., P.P., E.P., G.I.

Investigation: F.M., P.P., G.I.

Resources: G.I.

Data curation: A.A.D., P.P.

Writing: G.I.

Visualization: P.P., G.I.

Supervision: G.I.

Project administration: G.I.

Funding acquisition: G.I.

## DECLARATION OF INTERESTS

The authors declare no competing interests.

## METHODS

### Mice

All experimental manipulations were performed according to Italian legislation (DL 26/214, EU 63/2010, Ministero Della Sanità, Roma) and FELASA recommendations for the care and use of laboratory animals. Animal research protocols were reviewed and consented to by the Italian Ministry of Health.

We used 6-9 weeks old C57BL/6J mice of both sexes (Jackson Laboratory, RRID:IMSR_JAX:000664). Mice were co-housed with their littermates (2-4/cage) and maintained on a 12 hr/12 hr light/dark cycle with food and water ad libitum. Four C57BL/6 mice were used for the two odors/three concentrations (0.01, 0.1, and 1% v./v.) experiments; two C57BL/6 mice were used for the four odors/two concentrations (0.01 and 0.1% v./v.) experiment; four C57BL/6 mice were used for the odor identity experiments; three C57BL/6 mice were used for the artificial inhalation experiment; four C57BL/6 mice were used for the bulbectomy experiment; five C57BL/6 mice were used for the AEN neurectomy experiment.

### Surgical procedures

Animals were anesthetized with isoflurane (3% induction, 1.5% maintenance) and placed on a custom-made feedback-controlled heating pad. Pre-operative analgesia was induced through intramuscular injection of a bolus (4µl/g) of carprofen (Rimadyl, 0.05%)/dexamethasone (0.01%) and scalp infiltration of a tetracaine solution (0.05%). Post-operative analgesia was provided through carprofen diluted in the water bottle (Rimadyl, 134μl/100ml) following the procedure. Silicone-based eye ointment was applied on the eyes to protect the corneas during the surgery.

Anesthetized mice were mounted in a stereotaxic frame. The scalp was shaved with shaving cream and cleaned with isopropyl alcohol and iodopovidone. The skin and the periosteum from the lambdoid to the frontonasal sutures were removed, and muscles were partially detached from the skull to expose the occipital bone and the parietal ridges. To increase adherence of the implant to the skull, superficial grooves were drilled in the frontal, parietal, interparietal, and occipital bones. After leveling the skull yaw, pitch, and roll to obtain a flat skull configuration, two small reference crosses were scored using a scalpel at bregma and at the entry point of the Neuropixels probe (AP: 2.0/2.1 mm; ML: −2.0/−2.1 mm from bregma), and the incisions were filled with surgical ink and covered with UV-curable acrylic (Optibond). Finally, a titanium headplate was mounted onto the arm of the stereotaxic manipulator and attached to the skull with cyanoacrylate (Loctite 454) and dental cement (Paladur).

### Habituation to the experimental rig

Mice were habituated to head-fixation and the experimental rig starting five days before the experiment. Three habituation sessions of increasing duration (20, 40, and 60 min) were run over consecutive days. During the familiarization and the recording sessions, the mouse sat inside a 3-D printed black plastic tube, leaving exposed only the head. A custom-made polyether ether ketone (PEEK) nose cone was positioned in front of the mouse, loosely fitting the mouse’s snout. The nose cone was used to record the breathing signal and deliver the odorants during the experiments.

### Craniotomies

Mice were anesthetized again and placed in the stereotaxic manipulator two days before the recording. Two small craniotomies were opened: one for probe insertion (AP: 2.0/2.1 mm; ML: −2.0/−2.1 mm from bregma) and the other over the contralateral posterior parietal cortex for the ground electrode. The dura mater was not removed. Finally, the two craniotomies were filled with surgical silicone (Kwik-Cast, WPI).

### Tracheostomy

To probe PCx responses to the mechanical stimulation of the nasal cavity, we resorted to artificial ventilation. The electrophysiological recordings were performed immediately after the surgical preparation. For this procedure, mice were kept anesthetized throughout the surgical procedure and the recording with urethane (0.9 g/kg). Following a previously developed method^77^, we placed a cannula into the nasopharynx via the upper part of the trachea to allow the artificial suction of air through the nasal cavity; another cannula was placed in the lower end of the trachea to allow the natural gas exchange with the lungs.

### Ethmoidal nerve neurectomy

The anterior ethmoidal nerve was sectioned under isofluorane anesthesia before the recording session. The ocular bulb was laterally retracted, the upper region behind the eye was infiltrated with tetracaine solution (0.05%), and the nerve was cut by blunt dissection with fine forceps at the exit from the ethmoid foramen. The cut was verified post-mortem by visual inspection.

### Bulbectomy

A craniotomy was opened above the OBs. Next, a blunted needle connected to a vacuum system was used to aspirate both OBs while applying a steady flow of Ringer’s solution over the craniotomy until the cribriform plate was thoroughly cleaned and all the olfactory nerves were removed. Finally, the emptied cavity was filled with a sterile gelatin sponge (Gelfoam) and covered with dental cement.

### Stereotaxic probe insertion

All recordings were performed using Neuropixels 1.0 probes. The probe was mounted to a dovetail and affixed to a steel rod held by a micromanipulator (Luigs and Neumann). Before insertion, the back of the probe was coated with a solution of DiI (Thermofisher) using a paintbrush. Next, the silicone plug was removed from the craniotomies, and the probe was positioned above the recording site craniotomy (AP: 2.0/2.1 mm; ML: −2.0/−2.1 mm from bregma). Next, the probe was advanced through the dura and the cortex at approximately 5 µm/s until reaching the PCx (DV: 5.2/5.5 mm from bregma). The exact position depended on identifying a region with high firing rates and breathing-coupled activity. Then, the probe was retracted by 100 µm to allow the brain tissue to settle for 30 minutes before starting the recording. An Ag/AgCl electrode placed over the contralateral posterior parietal cortex was used for grounding. After electrode insertion, craniotomies were covered with a drop of agar solution (1% in Ringer’s solution). Neuropixels data were acquired and recorded at 30 kHz through a PIXe interface board connected to a PC.

### Experimental signals

Four signals were acquired besides the neural data: (1) a breathing signal generated by a flow sensor (AWM3300V Honeywell) plugged into the nose cone; (2) an odor signal generated by a photo-ionization detector (200B miniPID, Aurora Scientific) that sampled the odorant inside the nose cone; (3) a TTL signaling the start of each odor trial and (4) a TTL signal generated by a Bpod Analog Input module (Sanworks) at the start of each inhalation. To identify the onset of an inhalation on-line, the breathing signal was fed into the Bpod Analog Input module, and a threshold was set at the breathing signal zero-crossing before the inhalation peak. All signals were digitized and recorded at 30 kHz using an Intan RHD2000 board. Finally, to align the neural data with the other signals, a 1 Hz TTL clock signal (50% duty cycle) or a barcode was generated by an Arduino One board and recorded by the PIXe board and the Intan board.

### Odor delivery

Odorant stimuli were delivered using 5 seconds of odorant pulses. Odorants were delivered in 16 blocks (trials). The order of the odorants was randomized within each block. The inter-stimulus interval was randomly drawn from an exponential distribution (mean: 40 s, min-max: 30-50 s).

Odorants were delivered using a custom-made Arduino-controlled 13-valve olfactometer that delivers up to 12 odorants separately. The 13th valve was used to deliver a blank stimulus (no odor) between odor presentations. A vacuum continuously exhausted lingering odors. Odorants were contained in 15 ml vials partially filled with glass beads (Sigma-Aldrich). The headspace and the volume of odorant inside the vial were chosen to ensure that the amount of odor delivered within and across trials was steady as assessed by PID measurements. Each vial was separately connected to an olfactometer valve through a 3 in.-long PTFE tubing. A custom software controlled valve opening and closing, enabling switching between odor vials and blank vial. At the beginning of a trial, the opening of an odor valve was synchronized with the onset of inhalation, as detected by the Bpod Analog Input Module.

Two streams of carbon-filtered air (F1 and F2) were independently routed to the nose cone at 1 l/min. The F1 stream consisted in odor (F1O; 0.1 l/min) and carrier (F1C; 0.9 l/min) streams mixed. Upon opening an odorant valve, the F1O flow was routed through the open vial to a PEEK manifold using a 3 in.-long PTFE tubing. Inside the manifold, the F1O flow from the open vial was mixed with the F1C flow to obtain the F1 airflow. The outlet of the mixing manifold was connected through a 0.5-in.-long PTFE tubing to a final three-way valve (V1). The F2 stream was directly connected to a second three-way solenoid valve (V2). The V1 and V2 valve outlets converged in a final 0.5 in. PTFE tubing connected to the nose cone. The F1 and F2 airflows reached the odor cone during odor-ON and odor-OFF epochs, respectively. The V1 outlet was open during the odor-ON epoch, whereas the V2 outlet was open during the odor-OFF epoch. At the beginning of a trial, the olfactometer opened the odor vial valve to load the tubing with odorized air up to the final valve V1, which diverted the flow outside the nose cone. After 2 seconds, the onset of an inhalation triggered an “odor-ON” TTL that switched the outlet opening of the V1 and V2 valves such that only the odorized airflow F1 entered now the nose cone. After 5 seconds from the odor-ON TTL, the odor vial valve closed. After 10 seconds from the closing of the odorant vial valve, the V1 and V2 valves were switched to the odor-OFF configuration.

The odor panel included two odors (alpha-pinene and eucalyptol) delivered at three concentrations (0.01, 0.1, and 1% v./v.), three odors (limonene, p-cymene, and methyl butyrate) delivered at two concentrations (0.01 and 1% v./v.) and four odors (eugenol, dicyclohexyl disulfide, *p*-cymene, and methyl 2-furoate) delivered at a single concentration (0.01%). All odors were purchased from Sigma-Aldrich. To determine the dilution of each odor, a calibration curve was generated. To this aim, the signal generated by a photo-ionization detector (200B miniPID, Aurora) upon delivery of an odor at three dilutions (no dilution, 1:8, and 1:160 in mineral oil, Sigma-Aldrich; 10 presentations per dilution) was recorded. Then, an exponential curve was fitted to the miniPID traces to obtain the coefficient A for each dilution. Next, a second-order polynomial was fitted to the three A coefficients of each odor. Finally, the dilutions corresponding to 0.1%, 1%, and 10% v./v. of undiluted odor were extrapolated from the fitted polynomial. The carrier airflow further diluted the odor concentration by 10 to reach the final concentration of 0,01, 0.1, and 1 % v./v.

### Artificial mechanical stimulation of the nasal cavity

The cannula in the nasopharynx was attached to a computer-controlled solenoid valve which was connected to a vacuum line. The opening of the valve drew air inside the nasal cavity. The air was deodorized through a carbon filter before entering the nose cone. We applied 150-ms pulses of suction at 750-ms intervals. We tested five flow rates that span the range of estimated nasal flow rates in mice (50, 100, 150, 200, and 250 ml/min).

### Histology

Mice were deeply anesthetized with 1 g/kg urethane injection and intracardially perfused with 1% phosphate buffer solution (PBS) followed by 4% paraformaldehyde (PFA). The brain was dissected and immersed in 4% PFA for 24 h. Following fixation, the brain was washed with PBS and sectioned coronally (80 μm) with a vibratome. Sections were counter-stained with DAPI, and images were acquired with a fluorescence microscope at 10x magnification. The Allen CCF open-source toolbox^78^ was used to reconstruct the probe location in 3D, and Brainrender^79^ was used to visualize the probe position.

### Analysis of the breathing signal

The breathing signal was bandpass filtered between 0.5 to 50Hz using the MATLAB FMAToolbox (http://fmatoolbox.sourceforge.net/) and smoothed with a Gaussian kernel (standard deviation: 25 ms). Inhalation peaks were identified as the local minima of the respiratory signal. Inhalation peaks closer than 65 ms from the next peak were discarded. Inhalation onset and offset were identified as the breathing signal zero-crossings before and after the inhalation peak. Inhalation length was defined as the difference between the inhalation offset and onset times. The breathing cycle was calculated as the difference between two consecutive inhalation onsets. The negative inhalation slope was defined as the slope of the line between the nearest two points before the inhalation peak in the z-scored respiration signal with a value equal to 10% and 90% of the inhalation peak amplitude. Similarly, the positive inhalation slope was calculated as the slope of the line between the 90% to 10% of inhalation amplitude points in the z-scored respiration signal after the inhalation peak.

### Inhalation clustering

Inhalation waveforms were isolated by extracting the breathing signal within a 100 ms-long window centered on the inhalation peak. Each waveform was normalized by its 2-norm and considered as a point in a high-dimensional space. The dimensionality of the inhalation space was reduced by applying PCA using the *‘pca’* function in MATLAB (Mathworks®). The projection of the inhalation vector onto the *n*-dimensional subspace spanned by the *n* first PCs (where *n* is the minimum number of PCs that explain at least 98% of total variance) was calculated. The inhalation vectors in the *n*-dimensional space were clustered with a Gaussian Mixture Model using the *‘fitgmdist’* function in MATLAB (maximum number of iterations: 10,000, regularization value: 0.05, replicates: 15, full covariance matrix). The Calinski-Harabasz index was used to determine the optimal number of clusters. The centroid vector for each cluster was defined as the mean vector of all inhalation vectors within the cluster. To consolidate the GMM clustering, a Hierarchical Cluster (HC) dendrogram was applied to the centroid vectors using the *‘linkage’* function in MATLAB (distance metric: correlation). The HC dendrogram from an individual experiment usually resulted in two or three clusters. Visual inspection of the dendrograms suggested that two clusters were similar and stemmed from the same branch; thus, we merged the two clusters with the same parent. Finally, we assigned the clusters to the “slow” and “fast” inhalation types based on the mean of the inhalation lengths within each cluster, with “fast” inhalation assigned to the class with smaller inhalation lengths. The clustering algorithm was cross-validated on the inhalations from eight mice by fitting the GMM on a training set consisting of all the inhalations from seven mice and then using the fitted model to predict the labels of the inhalations from the held-out mouse. The ground-truth inhalation labels for the held-out mouse were obtained using the above clustering pipeline. This process was repeated for all possible combinations of test and train sets, and the classification accuracies were finally averaged. This clustering pipeline classified the inhalation label of held-out inhalations with 98% accuracy.

### Spike sorting and inclusion criteria

Spikes were sorted using KiloSort3^80^. All clusters were manually curated using Phy (https://github.com/cortex-lab/phy). During the manual curation process, the quality of every single unit assigned by the automatic sorting algorithm was evaluated based on the violation of the refractory period, the firing rate, and the stability of the firing rate during the recording session. Specifically, units that did not meet the quality criteria were removed from further analyses. Additionally, single units with a firing rate of less than 0.5 Hz were excluded from the subsequent analyses; this threshold was lowered to 0.1 Hz in the experiments with tracheostomized mice because the urethane anesthesia reduced the baseline firing rate.

Single units were further classified as regular (RS) or fast spiking (FS). A weighted average of the contributing cluster templates was computed for each unit, obtaining an average template waveform. Next, three features were extracted:

1. The latency between the negative and the following (post-depolarization) positive peak of the average waveform;
2. The asymmetry between the pre-depolarization (p1) and the post-depolarization (p2) peaks of the average waveform, computed as

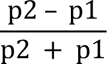
3. The average firing rate of the unit across the whole recording.

Finally, a k-means clustering (with Squared Euclidean distance, 10,000 iterations, and 2,500 replicates) of the features was used to partition the units recorded from each mouse into two categories. Units labeled as ‘RS’ had, on average, lower firing rates, larger trough-to-peak latency, and smaller asymmetry compared to units labeled as ‘FS’.

### Analysis of sniff-by-sniff responses

Inhalation and odor response were measured as the average spike count within a 70 or 180 ms window after the inhalation onset. Smoothed peri-event time histograms (PETH) were obtained by convolving the spike-times series within an inhalation with a Gaussian kernel (standard deviation: 10 ms), then averaging across all inhalations of the same type and finally subtracting the average firing rate across the odor-free epochs of the entire recording session. PETHs were aligned at the inhalation onset. Odorless-air inhalations were defined as those occurring within the −13 to −4 seconds window before the delivery of an odor. This window was considered sufficient to remove any lingering odor because the minimum time interval between two odor presentations was 30 s, and a vacuum line constantly exhausted odorants from the nose mask. Sniff-by-sniff odor responses were measured in a time window starting 400 ms after the odor onset and lasting for 4.6 s. The first two trials of each odor-concentration stimulus were excluded to avoid any bias due to the novelty of the stimulus. Furthermore, we excluded the first 400 ms of odor presentation because this period was dominated by fast sniffs (Figure S2D) and the typical transient response. Therefore, we did not analyze odor responses during the first inhalation of an odor.

### Concentration discrimination index

To measure how well a neuron could discriminate two concentrations, we first computed an area-under-the-receiver-operating-curve (auROC) using the spike counts during the inhalation of 0.01 and 0.1% v./v. odorant. The CDI was computed as abs(1 - 2*auROC). To generate the spike count distribution for each concentration, a random sample of inhalations was taken during the presentation of that concentration; an equal sample size was used for the two concentrations. Next, three different CDI were calculated based on a sample of slow inhalations, fast inhalations, and mixed slow and fast inhalations; 100 samples were drawn for each set of inhalations, and an average CDI was calculated for slow, fast, and mixed inhalations.

### Tuning curves

To obtain neural tuning curves during artificial stimulation of the nasal cavity, the average response for each airflow rate was calculated as the difference between the number of spikes fired by a neuron during the (0 500] and (−500 0] ms window where 0 is the onset of the air pulse.

To obtain neural tuning curves during natural breathing, inhalations were sorted into ten quantile bins based on the amplitude of the peak of airflow as measured by the flow sensor. This was done to ensure that each bin contained an equal number of inhalations. The tuning curve was generated by using the average response to an odorless inhalation during the first 180 ms for each bin.

Tuning curves were z-scored, and their slopes were calculated by linearly regressing the neural response to the airflow rate.

### Encoding models

A Poisson Generalized linear model (GLM) with regularization was employed to estimate the contribution of inhalation speed, odor concentration, and their interaction with the sniff-by-sniff odor response of a neuron. The model aimed to predict the spike count (*y_i_*) of neuron *i* during a single inhalation (either first 180 ms or 70 ms) using the following equation:

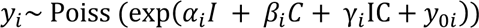

Here, the *I* term represents the inhalation speed (0 for slow and 1 for fast), and the *C* term represents the odor concentration (logarithm of the three concentrations 0.01, 0.1 and 1% v./v.).

*y*_0_*_i_* is a baseline bias of the response. The GLM was fitted using the *glmnet* toolbox (https://hastie.su.domains/glmnet_matlab/) in MATLAB (Mathworks®) with elastic net regularization, where the parameter *alpha* controlled elastic net penalty. A value of 0.95 was used for *alpha*, which smoothly interpolates the gap between lasso (*alpha* = 0) and ridge regression (*alpha* = 1). The optimal elastic net penalty value was selected using a 10-fold cross-validation approach.

We used a linear regression model for the simulations shown in Figure 5E-I and Figure 6B-C. In this case, the normalized firing rate *r_i_* of neuron *i* during the first 180 ms of inhalation was modeled with the following equation:

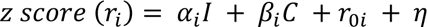

Here, the *I* term represents the inhalation speed (0 for slow and 1 for fast), and the *C* term represents the odor concentration (varying from 0 to 1 based on the logarithm of the concentration). The term *r*_0_ is a baseline bias, and η is a Gaussian noise term. To fit the models, inhalations were randomly sampled from all inhalations to have an equal size for each combination of inhalation speed and odor concentration term. The vector of the neural responses to the inhalations was z-scored and passed to the model with the corresponding design matrix. The significance of the model’s coefficients was assessed using an ANOVA test. The random sampling was repeated 100 times, and the coefficients of each term and their p-values were averaged across all re-samplings.

### Pseudo-population response matrix

The pseudo-population response matrix was obtained by pooling all recorded neurons’ sniff-by-sniff spike count responses to the inhalation events from all animals within the same experimental design. To create this matrix, an equal number of inhalation events for each odor concentration were randomly drawn from all inhalation events of all mice. Half of these events were fast inhalations, and the other half were slow inhalations. This allowed for an equal representation of both inhalation types in the sample. The same number of inhalations was sampled for each concentration, and then all inhalations were pooled. The response of each neuron to each inhalation in the selected sample was then vertically concatenated to obtain a population response vector during an inhalation. Finally, the response vectors of all neurons were concatenated horizontally to create a pseudo-population response matrix.

### PCA embeddings

To visualize the neural representation of odor concentrations across different inhalation speeds, PCA analysis was performed on the response covariance matrices obtained from a neural pseudo-population or single mouse neural populations. Before applying PCA, the neural responses of individual neurons were z-scored. PCA was also used to determine the projections onto the first 15 PCs of the sniff-by-sniff responses; these projections were separately used to decode inhalation speed in Figure 6B and Figure S7B,D and odor concentration and identity in Figure 6B,D and Figure S7B,D-F.

### Analysis of the angle between the concentration and the inhalation encoding axes

The encoding direction for odor concentrations and inhalation speed was determined using concentration-encoding unit vectors and sampling-encoding unit vectors; these two vectors were defined as unit vectors in the sniff-by-sniff response space. Linear discriminant analysis (LDA) was used to calculate the encoding vectors. Specifically, a pseudo-population response matrix with corresponding labels for each odor concentration and inhalation speed was used to fit the LDA model. This operation was performed using the Python *scikit-learn* library^81^. The eigenvalue decomposition solver was used to fit the LDA model with a shrinkage parameter that was automatically calculated by the Ledoit-Wolf lemma algorithm. The encoding vector was taken as the first column of the LDA transform scaling matrix. For each inhalation speed, the concentration-encoding unit vector was obtained from the LDA fit by using a pseudo-population response during a given inhalation speed and concentration using a different label for each concentration. Similarly, the inhalation-encoding unit vector was obtained from the LDA fit using pseudo-population response during either inhalation speed, and the type of inhalation speed was used as a label. Finally, the average angle between the concentration-encoding vectors for each inhalation speed and the average angle 0 between the concentration-encoding vectors and inhalation-encoding vectors were calculated and then transformed as: Θ’ = 90 - |90 – Θ|. This procedure was repeated 100 times with pseudo-population response matrices built using different randomly sampled neurons for each run.

### Inhalation speed decoding analysis

A linear support vector machine (lSVM) classifier was utilized for all classification analyses (inhalation speed, odor concentration, odor identity, and generalization analysis). The lSVM was implemented using the MATLAB (Mathworks®) Neural Decoding Toolbox (NDT)^82^ and *libsvm* (http://www.csie.ntu.edu.tw/~cjlin/libsvm/) toolbox. A 10-fold cross-validation was performed for all models. Before model fitting, the sniff-by-sniff neural responses in the training and test set were normalized by subtracting the means and standard deviations calculated from the responses in the training set.

To decode the inhalation speed, a pseudo-population response matrix was generated using only the spike counts during 180 ms windows for each inhalation of odorless air. Specifically, 400 random slow and fast inhalations were drawn, and the spike counts in 180 ms windows were used to build the response matrix of a pseudopopulation or an individual mouse population. The decoding accuracy for different numbers of neurons was determined by drawing random subsamples of neurons of different numerosity. This resampling process was repeated 100 times for each number of neurons to obtain reliable estimates of decoding accuracy. Additionally, the entire classification process was repeated 100 times, randomizing the sampled inhalation events included in each run. The reported accuracy for each number of neurons represents the mean of all resampling processes.

### Concentration decoding analysis

To classify odor concentration (0.01, 0.1, and 1% v./v.), a pseudo-population response matrix or a single animal population response matrix was built using the same number of inhalations for each odor-concentration pair to avoid any potential bias due to different inhalation sample sizes. Next, for each odor, corresponding rows of this matrix or its PC projections were sorted and passed to the classifier to decode the concentration in either the whole population response space or its PC projections space. This procedure was repeated 100 times with pseudo-population response matrices with different randomly sampled inhalations. The reported accuracies are the mean and standard deviation across all resampling processes.

To investigate the geometry of the neural space, a generalization paradigm was employed. To this end, a lSVM trained on data from one inhalation speed was tested on data from the other inhalation speed (*trans*-decoder). The average accuracy of *trans*-decoders was compared to that of *cis*-decoders that had been trained and tested instead on the odor responses during the same inhalation speed.

To assess the importance of heterogeneous selectivity for mechanosensory and olfactory inputs for sniff-invariant odor representations, we first fit the sniff-by-sniff responses of individual neurons with a linear regression model including a concentration (C = 0.01, 0.1, and 1%) and an inhalation speed (slow: I = 0; fast: I = 1) regressor. Then we selected a sub-pseudo-population with highly correlated mechanosensory (**α)** and olfactory regressor coefficients (**β**). To this end, a sub-pseudo-population of neurons meeting the following conditions were considered positively correlated: (|**β**| >= |**α**\*tan(pi/12)|) and (|**β**| <= |**α**\*tan(5*pi/12)|) with **α** and **β** having the same sign. Finally, another subset of neurons with **α** and **β** having different signs was selected to build a sub-pseudo-population with negatively correlated mechanosensory and concentration regressors. The union of these two sets was considered as the uncorrelated sub-pseudo-population. Then, an equal number of neurons was randomly sampled from these three sub-pseudo-populations, and their sniff-by-sniff odor responses were used by *cis*- and *trans*-decoders.

### Inhalation phase decoding

To predict the inhalation phase from neuronal activity, we employed Support Vector Regression (SVR) using the *scikit-learn* library^81^.

*Preprocessing.* First, the firing rate for every neuron was computed using a rolling gaussian window with sd = 20 ms (gaussian_filter() from scipy.ndimage). The firing rate matrix and the respiration trace were then downsampled to 10 ms bins to speed up the following computations. For every animal, the analysis was restricted to areas with at least 20 recorded neurons.

*Inhalation phase*. Analysis was restricted to the inhalation events defined and classified as described in the section *Inhalation clustering*. The inhalation phase was a number in the interval [0, 1] where 0 and 1 corresponded to the beginning and the end of the inhalation event, and the other values were linearly interpolated.

*PCA decomposition.* First, neuronal activity from all the neurons of each area was projected over principal components using the PCA class from sklearn.decomposition. To focus on the most relevant components for the prediction, PCs up to a cumulative explained (relative) variance > 0.5 were included (at least two PCs were always selected). We note that the analysis results hold for different inclusion criteria for the number of PCs.

*Epochs definition.* For the training and testing of the model, suitable non-overlapping epochs were created by concatenating inhalation periods for a total duration of 30 s. Depending on the inhalation number and classification, this resulted in a variable number of epochs (between 2 and 4) for every experimental animal. For every animal, all permutations of pairs of those suitable epochs were used as test and training datasets. Results were robust to different choices of epochs (i.e., more epochs with shorter windows).

*SVR training.* An SVR model was trained using SVR from sklearn.svm(), and its default parameters (kernel = ‘rbf’, degree = 3, gamma = ‘scale’, coef0 = 0.0, tol = 0.001, C = 1.0, epsilon = 0.1, shrinking = True; where ‘rbf’ kernel is the Radial Basis Function, and ‘scale’ gamma is 1 / (n_features * X.var())). The source data was the (n_components, timepoints) array of PC data; the variable to predict was the (timepoints,) array of inhalation phase data. The shuffle distributions were calculated by randomizing the inhalation phase array before prediction.

*Evaluation.* For every permutation of the epochs, the inhalation phase was predicted at every time point, and the performance of the model was assessed by calculating the squared Pearson correlation between the ground truth and the predicted data and then averaged across permutations.

### Decoding of concentration over time and phase bins

*Feature vectors generation.* First, we generated time- and phase-binned vectors of neuronal activity after the inhalation onset as follows:

- Time-binned vectors of 180 ms: the firing rate traces of all neurons in an area were used, as defined in the above section *Decoding of the inhalation phase. Preprocessing.* The traces were cropped from the onset of each inhalation in a 180 ms window and binned in 10-ms bins. Then, the traces of all neurons were concatenated to obtain, for every inhalation, an array with size (n_timebins x n_neurons), with n_timebins = 18.
- One-bin time vectors: the time-binned arrays defined above were averaged over the time bins.
- Time-binned vectors cropped over a full inhalation cycle: the respiratory phase was computed using the Hilbert transform. The time window in which the respiratory phase increased by 2π was found for every inhalation. Then, the firing rate trace was cropped in this window, padding it with zeros to account for the different durations of each inhalation period to a total of 500 ms (enough to accommodate > 99% of all inhalations). The arrays were then binned in 10-ms bins and concatenated across neurons to a total of (n_timebins x n_neurons) for every inhalation, with n_timebins = 50.
- Phase-binned vectors: the respiratory phase was computed using the Hilbert transform. The time window in which the respiratory phase increased by 2ρχ was found for every inhalation. The firing rate trace was cropped in this window and then binned into 36 bins equally spaced in phase (in this case, no zero-padding was necessary). Next, the activity of all neurons was concatenated to obtain an array with length (n_phasebins x n_neurons) for every inhalation, with n_phasebins = 36 and zero-padding at the end to ensure array length consistency.

*Odor concentration decoding.* To decode odor concentration from the firing rates, we used an SVM classifier using a generalization procedure like the one described above. First, inhalations were sampled 100 times with replacement for every animal to obtain a consistent number of events for each combination (inhalation_type, concentration). Then, ten-fold validation was used for training and testing the classifier for each extraction. Data were split into ten blocks, and the SVM trained over nine blocks and tested over the remaining one iterating over all ten possible left-out blocks. The SVM model was trained using SVC from sklearn.svm, and its default parameters but for the linearity of the kernel (kernel = linear, gamma = ‘scale’, coef0 = 0.0, tol = 0.001, C = 1.0, epsilon = 0.1, shrinking = True; where ‘scale’ gamma is 1/(n_features * X.var())). The source data was the ((n_bins x n_neurons), n_inhalations) array of concatenated firing rates over time for each inhalation; the variable to predict was the (n_inhalations) array of concentrations presented during each inhalation. For each iteration, the SVM was evaluated by computing the fraction of correct predictions. This number was then averaged across all folds and samplings (1000 different classifications) to obtain the numbers reported in the figures. Finally, the analysis was repeated separately for the two presented odors, and the results eventually merged.

### Simulation of neural responses with varying levels of inhalation-dependent concentration change

We tested whether integrating odor-independent mechanosensory inputs in the PCx code offsets the odor concentration alteration (ΔIC) inside the nasal cavity due to changes in inhalation speed. To this end, we compared odor concentration decoding using the activity pattern of two simulated populations of neurons with or without mechanosensory inputs. We reasoned that the ΔIC should be proportional to the external concentration and additive. Thus, we simulated the sniff-by-sniff responses of each neuron using the following Poisson model:

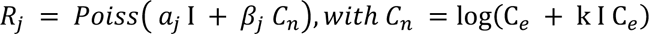

*R_j_* is the simulated neural response during a sniff. *α_j_* and *β_j_* are the mechanosensory and olfactory regressor coefficients previously estimated using a linear regression model fitted to the actual responses of neuron *j*. I is equal to 0 for slow inhalations and 1 for fast inhalations. C_e_ is the external odor concentration (0.01, 0.1, and 1%). k is a proportional factor that was parametrically changed in the range between 0 and 1 to simulate different levels of ΔIC; k = 0 means that a slow-to-fast change in inhalation speed does not change the odorant concentration (ΔIC = 0), whereas k = 1 means that a slow-to-fast change in inhalation speed increases the odor concentration inside the naris by 100% of the external concentration (ΔIC = C_e_). The population of neurons without mechanosensory inputs was generated by setting *α* to 0. *α* was set to 0 to simulate the population of neurons without mechanosensory inputs.

### Orthogonal representations in heterogeneous mixed-selectivity codes

Heterogeneous mixed selectivity of neural responses has an immediate connection with the orthogonality of population representations. This can be seen in a simple model, as follows. Consider a population code where the firing rate r_i_ of cell i is

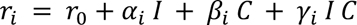

where, as above, I is the inhalation speed, and C is the concentration. For a population of size N (1≤i≤N), we can also write this in vector form as

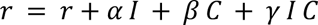

where r, α, β, and γ are now vectors with N entries. Assume that the number N of neurons is large and that coding is heterogeneous (that is, there is no special structure to the code), so that α, β, and γ are random vectors in N dimensions. In this case, α can be thought of as a scalar **|**α**|** controlling the overall intensity of tuning for inhalation speed in the population, times a random vector on the unit sphere, and the same for β and γ. The direction along which the population vector r encodes the concentration is

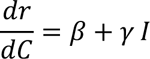

In other words, the concentration C is encoded in the direction β for slow inhalation (I=0) and the direction β + γ for fast inhalation (I=1). The cosine of the angle θ between the two encoding directions is

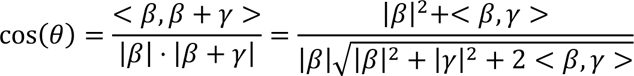

However, if β and γ are high-dimensional and their direction is chosen at random, it can be assumed they are approximately orthogonal^83^, and <β,γ> ≈0. Accordingly,

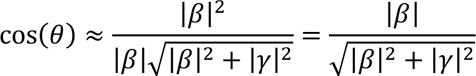

Therefore, the concentration encoding directions will tend towards orthogonality in the presence of strong interactions (when γ>> β, cos(θ)=0) and will be parallel when interactions are absent or weak (when |γ|=0, cos(θ)=1).

### Statistical tests

Sample sizes were not estimated in advance. Data groups were tested for normality using Kolmogorov-Smirnov test and then compared using the appropriate test. For regression modeling, confidence intervals were computed over bootstraps (with replacement) of the data. Statistical tests used, the value of n, and what n represents in each analysis can be found in the corresponding figure legend.

## SUPPLEMENTAL INFORMATION

**Figure S1.**
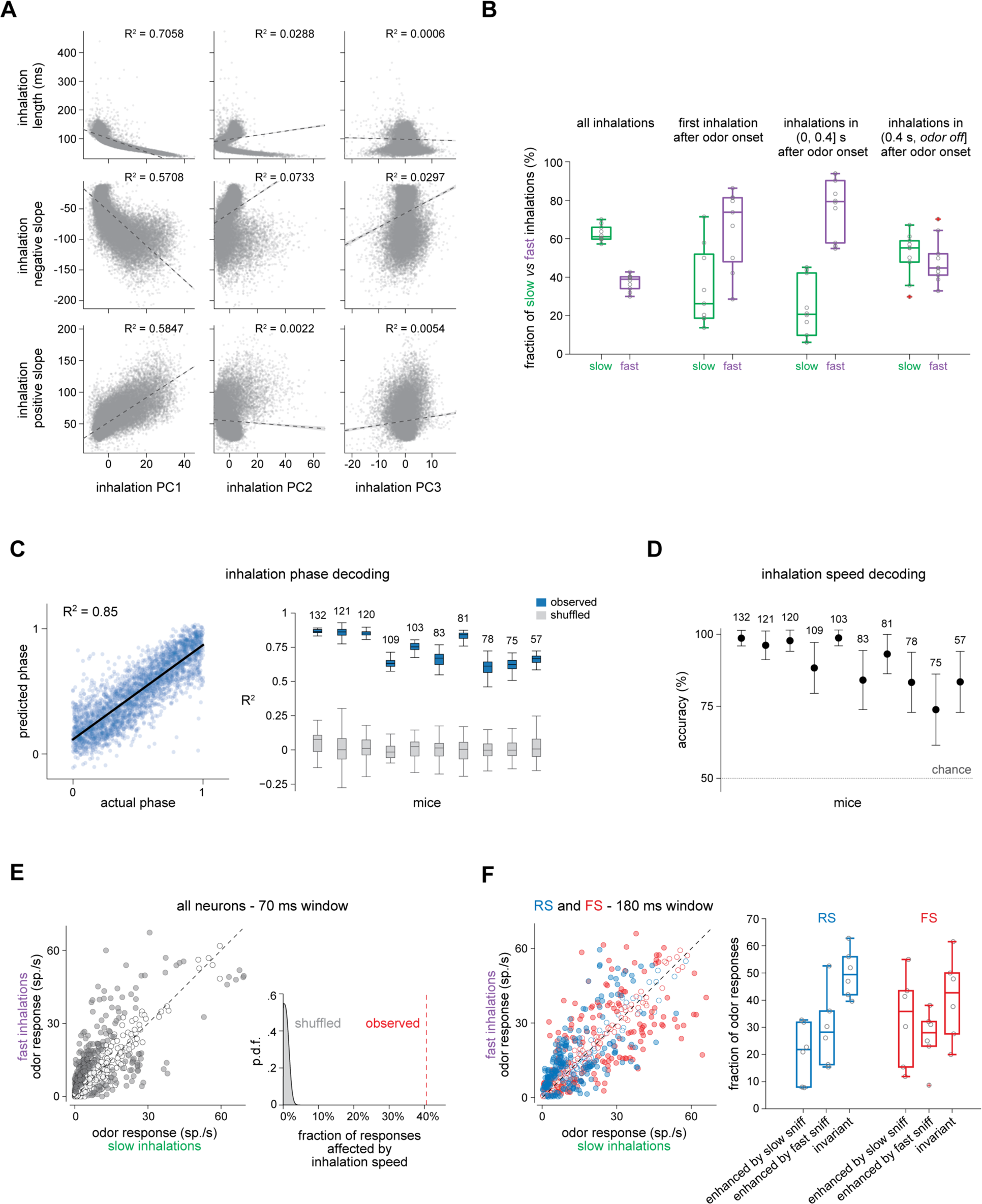
Sniffing causes unstable odor responses in the PCx. **(A)** Correlation between first (left), second (middle), and third (right) PCs and inhalation length (first row), descending slope (second row), and ascending slope (third row) of the inhalation waveform. Each data point corresponds to an inhalation. **(B)** Boxplots showing the fraction of slow and fast inhalations observed during the whole experiment, the first inhalation of an odor, the early 400 ms of the odor window, and the remainder of the odor window. Each data point represents a mean fraction from one experiment (N = 10 mice). Boxplots indicate the median, 25th and 75th percentile (box edges), and 1.5 times the interquartile range (whiskers). **(C)** Decoding the inhalation phase from the population activity in PCx. Left, scatterplot of actual and predicted inhalation phase for an example mouse (n = 120 neurons). The inhalation phase was predicted with a cross-validated support vector regression model. Right, goodness of prediction (R^2^) using the PCx neural trajectories in individual mice (blue). Gray, null distribution. The number above each boxplot indicates the number of neurons used for decoding. **(D)** Decoding the inhalation speed (fast *versus* slow) using the mean firing rate in the early 180 ms of inhalation in individual mice. Dotted line, chance level. The number above each boxplot indicates the number of neurons used for decoding. **(E)** Left: scatterplot of odor concentration responses during the early 70 ms window of inhalation (mean firing rate across slow or fast inhalations; five odors, 0.01 and 0.1% v./v. concentration; n = 687 responses) for all odor-responsive neurons (*P*-value < 0.01; n = 195 neurons, six mice). Gray dots, significant response difference between fast and slow inhalation; white dots, no statistical difference (rank-sum test, two-tails, *P*-value < 0.01). Right: fraction of neurons changing odor concentration response with inhalation speed (red vertical line) in the early 70 ms window and null distribution obtained by shuffling 1000 times the inhalation speed label for each odor response. **(F)** Left: scatterplot of odor concentration responses (n = 687 neuron-concentration pairs) after sorting neurons in RS (blue, n = 434 neuron-concentration pairs; 150 neurons) and FS (red, n = 291 neuron-concentration pairs; 94 neurons). Right: boxplots showing the fraction of odor concentration responses significantly enhanced by a fast or a slow inhalation and of those responses that were sniff-invariant (*P*-value < 0.01, rank-sum test) for RS and FS neurons. Each data point represents the fraction in a single experiment (N = 6 mice). Boxplots indicate the median, 25th and 75th percentile (box edges), and 1.5 times the interquartile range (whiskers).

**Figure S2.**
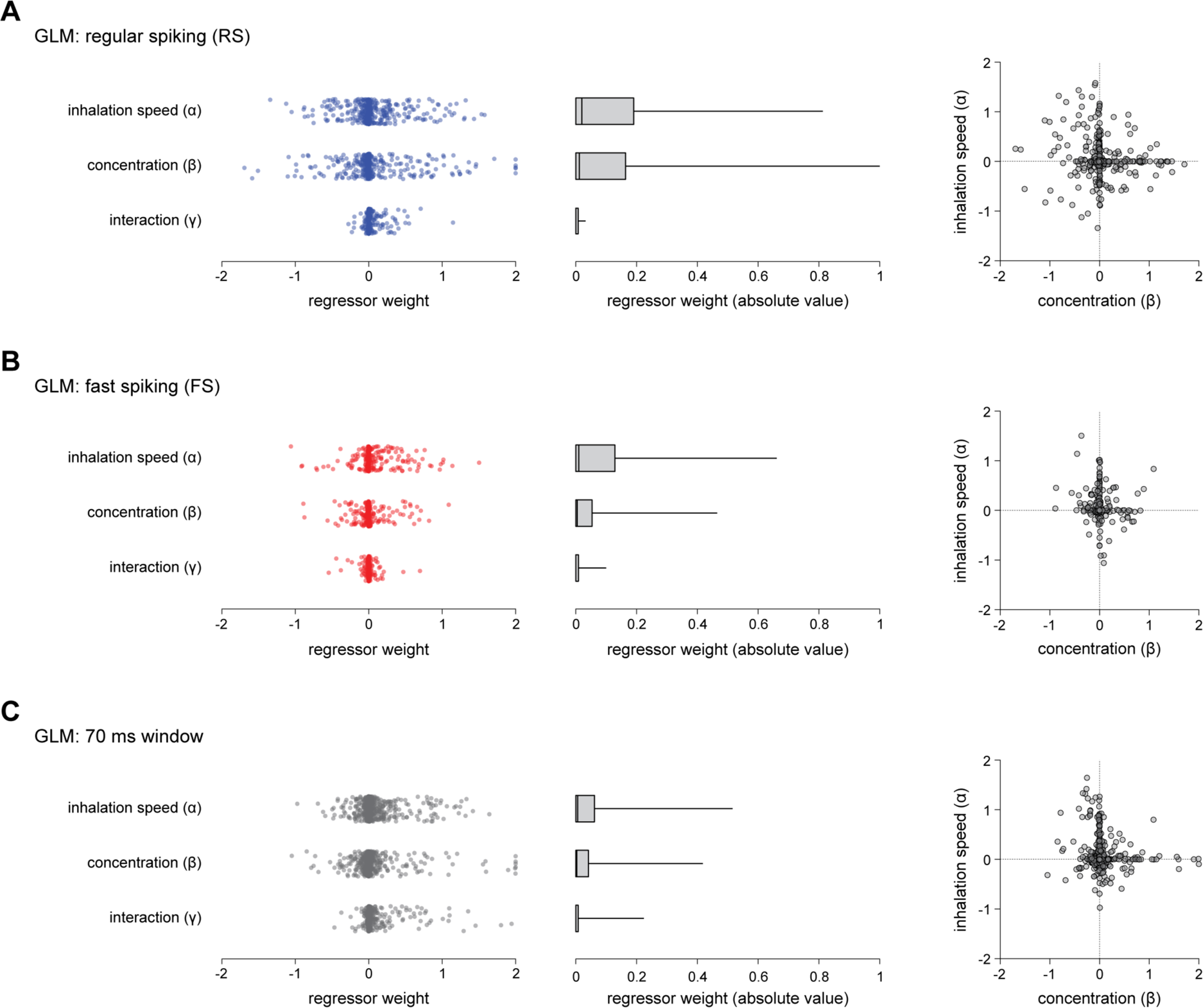
PCx neurons randomly mix olfactory and non-olfactory inputs during inhalations. **(A)** Left, distribution of the regularized GLM regressor weights for inhalation speed, odor concentration, and their interaction in RS neurons using the 0-180 ms window of an inhalation. Center, boxplot of the absolute values of the regressor weights indicating the median, 25th and 75th percentile (box edges), and the 95^th^ percentile (whisker). Right, scatterplot of the regressor weights for the concentration and inhalation speed factor. Each data point represents a neuron-odor pair (578 neuron-odor pairs; 289 neurons). **(B)** Same as (A) but for FS neurons (350 neuron-odor pairs; 175 neurons). **(C)** Left, distribution of the regularized GLM regressor weights for inhalation speed, odor concentration, and their interaction. Center, boxplot of the absolute values of the regressor weights indicating the median, 25th and 75th percentile (box edges), and the 95^th^ percentile (whisker). Right, scatterplot of the regularized regressor weights for the concentration and inhalation factor in all neurons using the 0-70 ms window of an inhalation. Each data point represents a neuron-odor pair (n = 928 neuron-odor pairs; 2 odors; 464 neurons; 4 mice).

**Figure S3.**
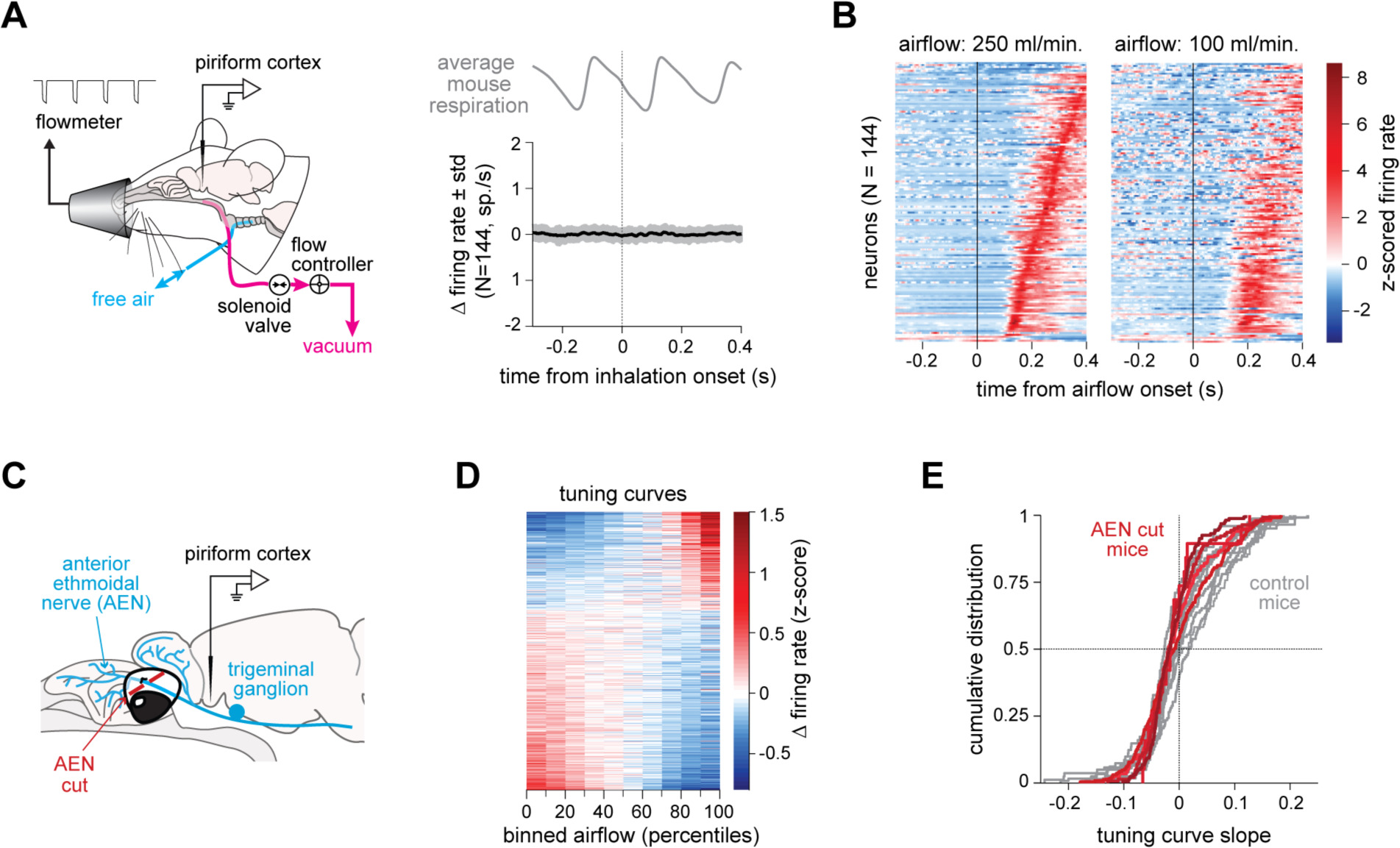
A mechanosensory input signals the nasal airflow rate to the PCx. **(A)** Left, experimental setup for artificial airflow stimulation of the nasal cavity in tracheostomized mice. Right, grand-average PETH aligned to the self-generated inhalation onset (n = 144 neurons responsive to nasal airflow stimulation, three mice). Self-generated inhalations were detected using a piezo sensor embedded in a chest belt. **(B)** Rastermaps of PETHs in tracheostomized mice. PETHs were aligned to the onset of a 150 ms-long airflow pulse applied to the back of the rhino-pharynx. Each row corresponds to the PETH of an individual neuron (n = 144 neurons, three mice). Left, PETH upon stimulation with a 100 ml/min airflow pulse; right, PETH upon stimulation with a 250 ml/min airflow pulse. PETHs were sorted bottom-up based on the latency to the peak of the response to the 250 ml/min airflow pulse. **(C)** Schematic of the AEN neurectomy experiment. **(D)** Raster map of peak airflow velocity tuning curves after severing the AEN (n = 551 velocity-tuned neurons of 752; 5 mice). **(E)** Cumulative distributions of the tuning curve slopes in control and neurectomized mice. Gray curves: control mice; red curves: neurectomized mice.

**Figure S4.**
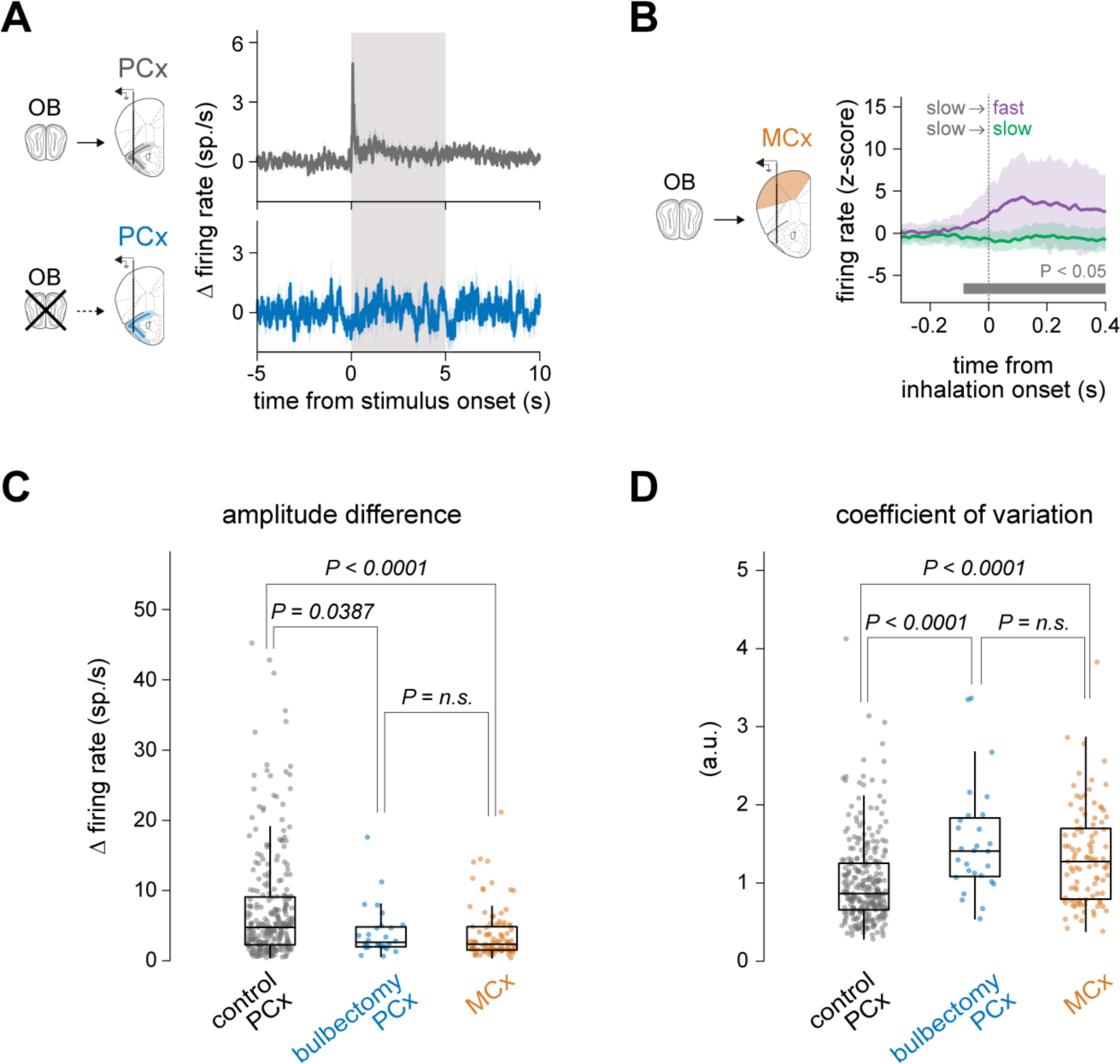
A top-down input supplies behavioral state information during a sniff. **(A)** Average odor response PETHs of all neurons in control (top; 660 neurons; six mice) and bulbectomized mice (bottom; 162 neurons; 4 mice). Shaded area, odor period (5 seconds). Unlike the other panels in this paper, here the PETHs are aligned to the onset of the first inhalation after the beginning of the odor presentation. **(B)** Average PETHs of motor cortex (MCx) neurons preferring a sniff in mice with intact OBs (99 of 548 neurons). Shaded area, mean ± s.e.m. The bar below the PETH indicates when the sniff responses are significantly bigger than the slow inhalation responses (Benjamini-Hochberg adjusted *P*-value < 0.05, one-sided t-test). **(C)** Difference between the amplitudes of the responses to the first sniff and a regular inhalation in the PCx of control and bulbectomized mice and in the MCx of control mice (P-value < 0.0001, 1-way ANOVA; Tukey-Kramer post hoc *P*-values shown). Boxplots indicate the median, 25th and 75th percentile (box edges), and 1.5 times the interquartile range (whiskers). **(D)** Coefficient of variation of the response amplitude across first sniffs in the PCx of control and bulbectomized mice and in the MCx of control mice (*P*-value < 0.0001, 1-way ANOVA; Tukey-Kramer post hoc *P*-values shown). Boxplots indicate the median, 25th and 75th percentile (box edges), and 1.5 times the interquartile range (whiskers).

**Figure S5.**
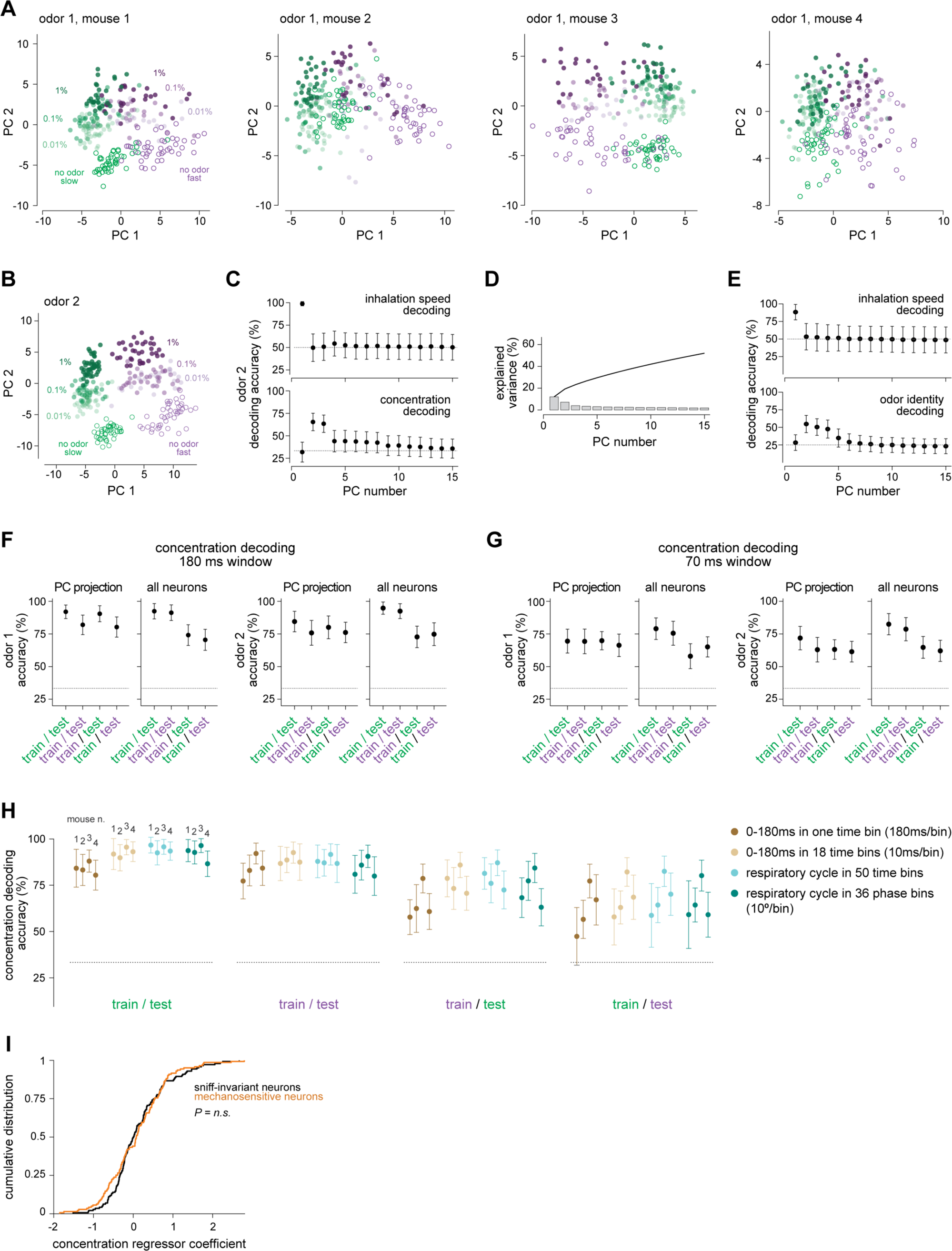
Odor and inhalation information are independent. **(A)** PCA embeddings of the inhalation-by-inhalation population odor responses to three concentrations of the same odor of Figure 5A (eucalyptol), but for individual mice. **(B)** PCA embedding of the inhalation-by-inhalation odor responses of a pseudopopulation of PCx neurons like in Figure 5A but for another example odor (pinene). **(C)** Decoding accuracies of inhalation speed and concentration using different PCs of a pseudo-population of PCx neurons like in Figure 5B but for the other example odor. **(D)** Scree plot of the variance explained by the first 15 PCs of the covariance matrix, including the responses of 464 neurons to the two example odors presented at three concentrations. **(E)** Decoding accuracy of inhalation speed (top) and odor identity (bottom) using the projection of the pseudo-population odor responses on each of the first fifteen PCs (four odors; two inhalation speeds). Mean ± standard deviation is shown. The data used for this analysis are from a different set of experiments where four odors were presented to mice without changes in concentration. **(F)** Concentration cis- and trans-decoding accuracy using PCx activity during the 0-180 ms window of inhalation. Mean ± standard deviation is shown. **(G)** Same as in (F) but using the 0-70 ms window of inhalation. **(H)** Comparison of the accuracy of a linear decoder across four different coding schemes. Dark brown: average population activity in the 0-180 ms window of a respiratory cycle; 0 ms: inhalation onset. Light brown: neural population trajectory during the 0-180 ms window; the neural activity during the 180 ms window was binned in 18 bins of equal duration (10 ms), and a population vector was built using the average activity of each neuron during a given time bin; finally, the 18 vectors were concatenated to obtain the population trajectory. Light teal: neural population trajectory during the entire respiratory cycle; the respiratory cycle was binned in 50 time-bins of equal duration within the cycle; the time bin duration varied across respiratory cycles due to the different duration of each cycle; the 50 population vectors were concatenated to obtain the population vector during a respiratory cycle. Dark teal: neural population trajectory during the entire respiratory cycle using the phase instead of the timing; the respiratory cycle was binned in 36 phase-bins of equal length (10°); the 36 population vectors were concatenated to obtain the population trajectory during a respiratory cycle. Mean ± standard deviation is shown for four mice. (I) Related to Figure 5G-I. Cumulative distribution of concentration regressor coefficients for mechano-sensitive and sniff-invariant neurons (Kolmogorov-Smirnov test, P-value = 0.4793).

## Notes

### Competing Interest Statement

The authors have declared no competing interest.

### Summary of Updates

All analyses were improved.

